# A pectin-binding peptide with a structural and signaling role in the assembly of the plant cell wall

**DOI:** 10.1101/2023.08.23.554416

**Authors:** Sébastjen Schoenaers, Hyun Kyung Lee, Martine Gonneau, Elvina Faucher, Thomas Levasseur, Elodie Akary, Naomi Claeijs, Steven Moussu, Caroline Broyart, Daria Balcerowicz, Hamada AbdElgawad, Andrea Bassi, Daniel Santa Cruz Damineli, Alex Costa, José A Feijó, Celine Moreau, Estelle Bonnin, Bernard Cathala, Julia Santiago, Herman Höfte, Kris Vissenberg

## Abstract

Pressurized cells with strong walls make up the hydrostatic skeleton of plants. Assembly and expansion of such stressed walls depend on a family of secreted RAPID ALKALINIZATION FACTOR (RALF) peptides which, curiously, bind both a membrane receptor complex and wall-localized LEUCINE-RICH REPEAT EXTENSINs (LRXs) in a mutually exclusive way. Here we show that, in root hairs, the RALF22 peptide has a dual structural and signaling role in cell expansion. Together with LRX1, it directs the compaction of charged pectin polymers at the root hair tip into periodic circumferential rings. Free RALF22 induces the formation of a complex with LORELEI-LIKE-GPI-ANCHORED PROTEIN 1 (LLG1) and FERONIA (FER), triggering adaptive cellular responses. These findings show how a peptide simultaneously functions as a structural component organizing cell wall architecture and as a signaling molecule that regulates this process. This mechanism may also underlie wall assembly and expansion in other plant cell types.

## Introduction

The pectic polysaccharides homogalacturonans (HGs), like glycosaminoglycans (GAGs) in the animal extracellular matrix (ECM)^1^, are abundant unbranched and charged polymers that play a critical role in the control of the physico-chemical properties of the plant cell wall (CW)^2^. These properties include the remarkable ability to expand while simultaneously resisting the tensile forces imposed by the turgor pressure on the CW^2^.

Pectins are galacturonic acid-containing polymers, which, together with hemicelluloses (e.g. xyloglucans) and (glyco)proteins, form the matrix that surrounds and connects cellulose microfibrils in growing cells^3^. HGs form the most abundant class of pectins, reaching up to 50% of the total CW polymer content^4^. They are synthesized in a highly methylesterified, uncharged form, and can be demethylesterified by wall-associated pectin methylesterases (PMEs), thus generating random or block-wise anionic charge patterns^2^. *In vivo* studies have associated HG demethylesterification with either the cessation or promotion of cell expansion, depending on the context^5^. Calcium (Ca^2+^)-mediated crosslinking of polyanionic HG has been proposed as a driver for CW stiffening and growth restriction^6,7^. Growth promotion instead, has been proposed to occur upon the exchange of load-bearing Ca^2+^ crosslinks with newly generated pectate, pectin swelling, or more complex scenarios involving HG turnover or feedback signaling^8–15^. In any case, the physicochemical mechanisms in the pectic CW underlying growth changes remain poorly understood so far.

In this context, it is interesting to note that the properties of the ECM in animals depend not only on the intrinsic physical properties of the GAG polymers but also on the interaction of GAGs and their core proteins with specific GAG binding proteins that modulate their structure^16^. For instance, whereas GAGs and GAG-rich proteoglycans promote ECM swelling and generate ultrasoft matrices, the presence of crosslinking proteins leads to water loss, compaction and rigidification of the same matrices^16^. In addition, compaction can lead to the transition from a homogenous phase to segregated microphases, as observed for instance for the formation of perineural nets, the porous ECM that wraps neurons^16^.

In plants, some *in vitro* evidence shows the interaction of pectin with basic versions of extensins, which are hydroxyproline-rich structural proteins that can form a crosslinked network in the CW. While *in vivo* confirmation is still lacking, these interactions are believed to be partly established by intermolecular di-isodityrosine oxidative crosslinks^17–20^. Despite the similarities between plant HG and animal GAG polymers, so far it is not known whether pectin-binding proteins contribute to the modulation of CW properties in plants.

Here we show that RALF22, a member of the Rapid Alkalinisation Factor (RALF) family is a driver for the structuration of the CW through its interaction with polyanionic pectin. RALF peptides were first identified by their capacity to induce a rapid alkalinization of the growth medium and are part of a family of 37 members in *Arabidopsis thaliana*^21,22^. The well-studied peptide RALF23 binds to the GPI-anchored protein LORELEI-LIKE-GPI-ANCHORED PROTEIN 1 (LLG1), thus triggering the formation of a ternary complex with the *Catharanthus roseus* Receptor-Like Kinase1-Like (CrRLK1L) protein FERONIA (FER), involved in immunity signaling^23^. RALF4 and 19 are essential for the growth of pollen tubes, in which they activate a module consisting of LORELEI-LIKE-GPI-ANCHORED PROTEIN 2 (LLG2), the CrRLK1L proteins BUDDHAS PAPER SEAL 1 (BUPS1) and ANXUR (ANX1), the cytosolic kinase MARIS and the Ca^2+^ channel of the MILDEW RESISTANCE LOCUS O (MLO) family^24–27^. Related pathways are required for sperm release from pollen tubes, CW integrity control, stress responses or mechanosensing^28^. Interestingly, several RALF peptides also tightly bind CW proteins of the LEUCINE-RICH REPEAT EXTENSIN (LRX) family, with distinct and mutually exclusive binding modes to the LLG-CrRLK1L signaling system^27,29–31^. LRXs are dimeric CW proteins, essential for growth, with a RALF-binding LRR domain and a C-terminal extensin domain rich in crosslinkable tyrosines, but without patches of basic residues that are found in pectin-binding extensins^29,32^.

In this study, we used *Arabidopsis* root hairs (RHs) as a model system to study RALF-mediated pectin structuration and the biological function underlying the existence of two distinct classes of RALF-binding proteins. RHs show highly polarized apical growth that can display oscillations in growth rate and a number of other parameters, such as cytosolic Ca^2+^ levels, CW pectin charge, Reactive Oxygen Species (ROS) levels and cytosolic and apoplastic pH^33^. This suggests that RH growth is regulated by processes that require, or are optimized by temporal synchronization, eventually leading to the emergence of periodicity in CW modifying activities. For instance, processive pectin demethylesterification is favored at neutral pH, whereas the remodeling of the cellulose-xyloglucan network by the CW-loosening expansins occurs at low pH^34,35^.

We show that a periodic CW architecture emerges in RHs from the charge-dependent compaction of polyanionic pectin through its binding to the CW protein complexes LRX1/2-RALF22. In the absence of RALF22 or LRX1/2, the CW architecture is severely perturbed, leading to a frequent loss of cell integrity. Furthermore, when not incorporated into the CW, the same peptide has a signaling role through the formation of a membrane ternary complex with LLG1-FER, which induces cellular changes that influence CW assembly and growth.

## Results

### RALF22 is required for root hair growth

To study RALF-regulated cell expansion, we screened publicly available transcriptome data for RH-expressed *RALF(s)*. From the seven *RALFs* that had the highest expression in the primary root (Fig. S1A) only *RALF22* transcripts were enriched in RH cells, as shown by single cell RNAseq data (Fig. S1B-C), and its expression correlated with the trichoblast developmental stage (Fig. S1D)^36^. Using CRISPR-Cas9, we generated a *RALF22* loss-of-function line by a 182-nucleotides deletion 51 nucleotides downstream of the start codon (*ralf22-1*; Fig. 1A). A second loss-of-function allele was identified from the GabiKat seed repository (*ralf22-2*; GK_293H09; Fig. 1A). Both lines displayed a distinct short and bulged RH phenotype with frequent cell bursting, which was fully complemented by re-introducing the wild-type *pRALF22::RALF22* sequence into the mutant genome (Fig. 1B-C; movie 1). Transcriptional reporter lines (Col-0 x *pRALF22::GFP*) accumulated GFP in RH cells only (Fig. 1D), and showed that *RALF22* transcription commenced during RH bulge formation, peaked during tip growth and ceased upon RH maturation (Fig. 1D). These data show that RALF22 is essential for normal RH morphogenesis.

**Figure 1.**
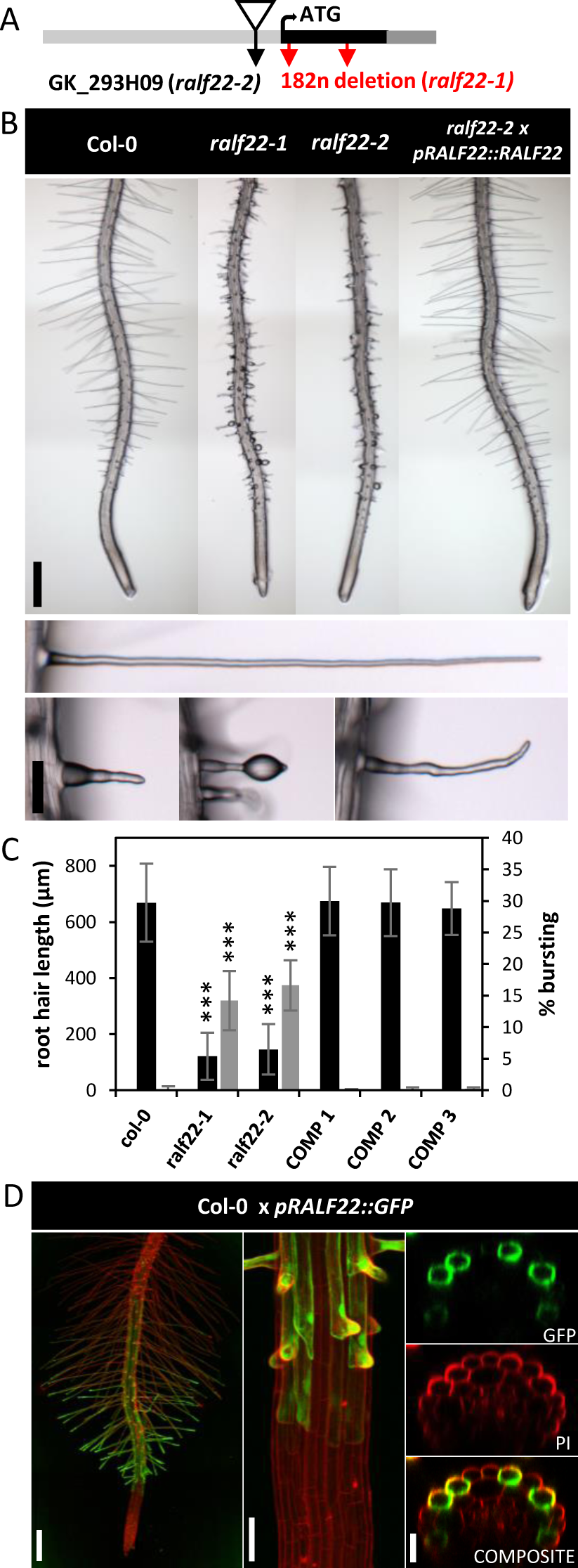
RALF22 regulates root hair growth. (A) scheme of the RALF22 genomic sequence showing the T-DNA insertion site of GabiKat line GK_293H03 *(ralf22-2)* and the 182 nucleotide-long deletion in *ralf22-1*. (B) representative 6-day-old roots of Col-0, *ralf22-1* (-/-), *ralf22-2* (-/-) and *ralf22-2 x pRALF22::RALF22* (COMP) seedlings (scale bar=500µm). Close-ups of RHs showing the short, bulged and burst (cfr. movie 1) *RALF22* loss-of-function phenotype (scale bar=100µm). (C) graph showing the mean RH length (±SEM) and % of RH bursting for all genotypes (n≥5, ***p<0.001). Three independent complementation lines (*ralf22-2 x pRALF22:RALF22*) are shown. (D) representative confocal maximal projections and transverse optical sections of 6-day-old Col-0 seedlings expressing GFP under the control of the RALF22 promoter (Col-0 x *pRALF22::GFP*). *RALF22* is transcribed in trichoblast cells throughout RH growth (scale bars= left, 100µm; middle, 50µm; right; 25µm).

### RALF22 has FERONIA-dependent and FERONIA-independent effects on root hair growth

To study RALF22 function, we investigated whether RALF22 is a RH-specific *Cr*RLK1L ligand, similar to other *Cr*RLK1L-binding RALF peptides^24,37,38^. Like other RALFs, RALF22 contains a propeptide sequence that is cleaved off by subtilisin-like serine proteases during protein maturation^39,40^. The mature RALF22 protein harbors two disulfide cysteine-bridges and a conserved YISY motif that is critical for receptor binding (Fig. 2A)^23,39^. To date, two *Cr*RLK1L receptor-like kinases (ERU, FER) and the FER-RALF23 co-receptor LLG1, were shown to be required for RH morphogenesis^41–43^. We recombinantly expressed LLG1 and the ERU and FER ectodomains (ERU_ecd_, FER_ecd_) in insect cells and purified them for interaction analysis (Fig. S2A-B). In addition, we chemically synthesized RALF22. We then quantified the binding affinities amongst all possible protein combinations (Fig. S2C) using microscale thermophoresis (MST). While RALF22 interacted with LLG1 with a dissociation constant (Kd) of 6.09±1.13µM (Fig. 2B), it did not directly interact with ERU_ecd_ or FER_ecd_ alone (Fig. S2C). Preincubated LLG1-RALF22, however, formed a high affinity (118.02±59.01nM) ternary complex with FER_ecd_, but not with ERU_ecd_ (Fig. 2C-D, Fig. S2C). Next, we substituted both tyrosines with alanines in the conserved RALF22 YISY motif (RALF22^Y75A,Y78A^) that mediates receptor binding^23^. Compared to wild type RALF22, we observed a 4.7-fold decrease in the affinity of RALF22^Y75A,Y78A^ for LLG1 (28.48±2.43µM) (Fig. 2B). In addition, LLG1-RALF22^Y75A,Y78A^ no longer formed a ternary complex with FER_ecd_ (Fig. 2C). Together, these results show that RALF22, like RALF23^23^, binds to LLG1 and nucleates the formation of a ternary LLG1-RALF22-FER_ecd_ complex.

**Figure 2.**
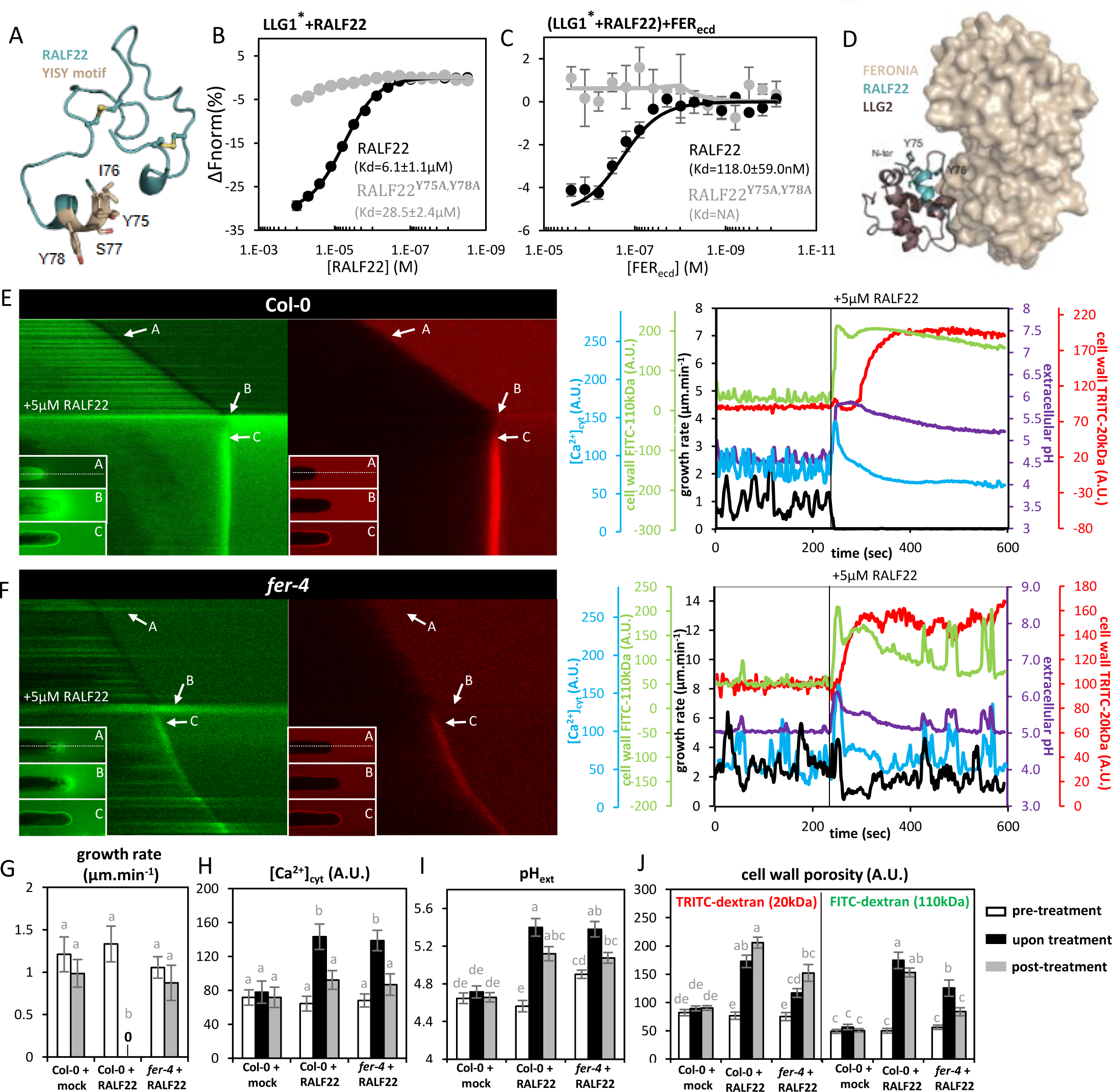

RALF22 binds to FERONIA to regulate root hair growth. (A) model of mature RALF22 showing the conserved YISY domain (sticks) required for LLG binding. (B) MST results showing the binding affinity between fluorescently labelled recombinant LLG1 and RALF22 (n=7) or RALF22^Y75A,Y78A^ (n=3). (C) Binding affinity between the FERONIA ectodomain (FER_ecd_) and fluorescently labelled LLG1 preincubated with RALF22 (n=3) or RALF22^Y75A,Y78A^ (n=3). MST values are represented as the average normalized change in fluorescence (%) ± SEM. (D) Homology model of the LLG2-RALF22-FER_ecd_complex. The N-terminal region of RALF22 was docked into the LLG2 structure (PDB:6A5E). FER_ecd_ was superimposed based on the crystal structure. The YISY motif is highlighted in sticks. (E-F) Timelapse imaging of Col-0 (E) and *fer-4* (F) RHs being treated with 5µM RALF22 (cfr. Movie 2). The kymographs and corresponding plots depict changes in the growth rate, [Ca^2+^]_cyt_ (cytosolic GCaMP3 fluorescence intensity), pH_ext_ (extracellular FITC-110kDa dextran/TRITC-20kDa dextran fluorescence intensity ratio) and migration of fluorescent FITC-110kDa dextran and TRITC-20kDa dextran into the CW (a proxy for altered CW physico-chemistry) during 4min of steady state growth followed by treatment with RALF 22. The RH’s response to RALF22 was monitored for 6min. Kymographs were generated along the horizontal lines depicted in the insets. (G-J) quantification of the average growth rate (G), [Ca^2+^] (H), pH_ext_ (I), and FITC or TRITC fluorescence in the CW (J) before, upon and after treatment with RALF22 in Col-0 (n=9) and *fer-4* (n=8) RHs. Col-0 RHs were also treated with a mock solution (-RALF22, n=7). Barplots represent the mean ± SEM. Different letters represent statistical significance (α=0.05).

Next, we explored the *in vivo* relevance of this interaction for RH growth. We optimized a protocol that allowed us to reproducibly treat growing RHs while imaging key parameters related to RH elongation, namely oscillations in growth rate, extracellular pH (pH_ext_) and intracellular calcium ([Ca^2+^]). To this end, we grew Col-0 and FERONIA loss-of-function (*fer-4*) seedlings expressing the cytosolic Ca^2+^ indicator GCaMP3^44^ in microfluidic chips in the presence of FITC (pH-sensitive) and TRITC (pH-insensitive) coupled to 110kDa and 20kDa neutral dextran, respectively. In animal systems, FITC or TRITC coupled to dextrans of different sizes are commonly used for permeability studies^45,46^. Here, the large dextran molecules restrict both fluorophores to the extracellular medium and the FITC/TRITC ratio provides a ratiometric quantification of the pH_ext_ (Fig. S3A). Using confocal microscopy, we monitored Col-0 and *fer-4* RHs before and after administration of 5µM RALF22. During steady-state growth, both Col-0 and *fer-4* RHs displayed oscillations in growth rate, [Ca^2+^] and pH (Fig. 2E, F). Upon RALF22 treatment, Col-0 RHs immediately stopped growing (Fig. 2E, G). This growth arrest was accompanied by a transient spike in [Ca^2+^] (Fig. 2E, H) and an increase in pH (Fig. 2E, I), which rapidly spread from tip to shank (Fig. S3B-D). The pH remained high over the next 6 min (Fig. 2E, movie 2). Surprisingly, RALF22 treatment induced a clear migration and accumulation of both dextran-coupled TRITC and FITC fluorophores into the CW, at a comparable rate, suggesting a change in the physicochemical properties of the CW (Fig. 2E, J, movie 2). Upon RALF22 treatment, *fer-4* RHs showed a mild growth inhibition and an increase in [Ca^2+^] and pH (Fig. 2F-I). In striking contrast to Col-0 RHs the extracellular alkalinization did not propagate towards the shank (Fig. S3B, E), while growth rate, [Ca^2+^] and pH oscillations recovered rapidly (Fig. 2F-I, movie 2). Crucially, the RALF22-induced migration and accumulation of dextran-coupled TRITC and FITC into the CW was still apparent in *fer-4* RHs (Fig. 2F, J; movie 2), suggesting that besides its FER-dependent signaling role, RALF22 treatment has an additional FER-independent effect on the physicochemical properties of the CW.

### RALF22 interacts with demethylesterified homogalacturonan and induces its compaction

To understand this FER-independent effect of RALF22 on the CW, it is interesting to note that mature RALF22 is a positively charged protein (isoelectric point = 10.53) (Fig. 3A), which potentially interacts electrostatically with poly-anionic demethylesterified homogalacturonan (HG) in the CW. We used MST to study the interaction between RALF22 and a fully demethylesterified oligogalacturonide, with a degree of polymerization between 7 and 13 (OG7-13)^47^. We observed a robust interaction with a Kd of 3.03±0.53µM (Fig. 3B). The charge dependency of the interaction is suggested by the 3.7-fold reduction in affinity (Kd = 11.1±2.65µM) for a mutant peptide RALF22^R82A,R90A,R100A^ in which 3 charged arginines were replaced by neutral alanine residues (Fig. 3B). To further explore this interaction, we used Quartz Crystal Microbalance with Dissipation monitoring (QCM-D). This technique measures the mass and viscoelasticity of polymer layers deposited on gold-coated oscillating quartz crystals, where a decrease in oscillation frequency (ΔF) linearly correlates with a mass increase and a decrease in the dissipation time of the oscillation after interruption of the current (ΔD) correlates with an increase in viscoelasticity of the layer (Fig. 3C)^48^. In our experimental setup, we first created a pectin layer on the quartz surface (Fig. S4A, B). To this end, the gold-coated quartz, which previously had been spin-coated with a positively charged polymer (poly-allylamine hydrochloride, PAH), was placed in a flow cell and a pectin solution was flown over the surface with a constant flow rate, while monitoring ΔF and ΔD (Fig. S4A, B). As expected, an elastic pectin layer was formed as shown by the decrease in ΔF_pectin_ and increase in ΔD_pectin_, both for pectin with a degree of methylesterification (DM) of 75% (DM75, Fig. S4A) and 31% (DM31, Fig. S4B; see Fig. S4C, D for a detailed characterization of the pectin preparations). After a washing step, RALF22 was applied to the DM75 pectin layer (Fig. 3D, Fig. S4A). Unexpectedly, this showed a mass decrease (seen as an increase in ΔF_RALF22_ corresponding to a 9.60±2.22% loss of the mass of the hydrated pectin layer, n=14) and a concomitant increase in stiffness (seen as a decrease in ΔD_RALF22_ corresponding to a 34.05±6.63% decrease in viscoelasticity of the pectin layer, n=14) (Fig. 3D, G). These results indicate that RALF22 interacts with pectin, but also induces water loss from the pectin layer, the mass of which exceeds the gain in mass due the binding of the 5.6 kDa RALF22 peptide. This causes stiffening of the layer, which is expected for the formation of a polyelectrolyte complex where the neutralization of the charges causes dewatering^49,50^. This interaction was stable under the experimental conditions used since no important changes in ΔF and ΔD were observed during the subsequent washing step (Fig. 3D, Fig. S4A). The charge dependence of the interaction was confirmed by the increased dewatering capacity of RALF22 on pectin with a lower degree of methylesterification (DM31; increase in ΔF_RALF22_ of 38.98±4.27%, decrease in ΔD_RALF22_ of 94.13±8.36%, n=15) (Fig. 3E, G, Fig. S4B) as well as the strongly reduced dewatering capacity of mutant peptide RALF22^R82A,R90A,R100A^ (increase in ΔF of 3.53±0.24%, decrease in ΔD of 13.11±1.64%, n=2) relative to wild-type RALF22 (Fig. 3F, G). The interaction with fully methylesterified HG could not be assessed using this technique since such polymers do not bind the PAH layer. Together, these results show that, *in vitro*, RALF22 binds and dewaters pectin in a charge-dependent manner. The formation of such a polyelectrolyte complex not only creates denser zones in the gel, but also augments the porosity of the non-compacted zones^16,49,50^. An analogous process is thought to underlie the formation of perineural nets, the porous aggregated matrix wrapped around the surface of neurons, which is formed through the interaction between charged GAGs and crosslinking proteins^16^. Similar RALF22-induced physicochemical changes (pectin charge neutralization, water loss and porosity increase) could account for the FER-independent accumulation of FITC- and TRITC-dextran in the CW of growing RHs (Fig. 2E, F, J; movie 2).

**Figure 3.**
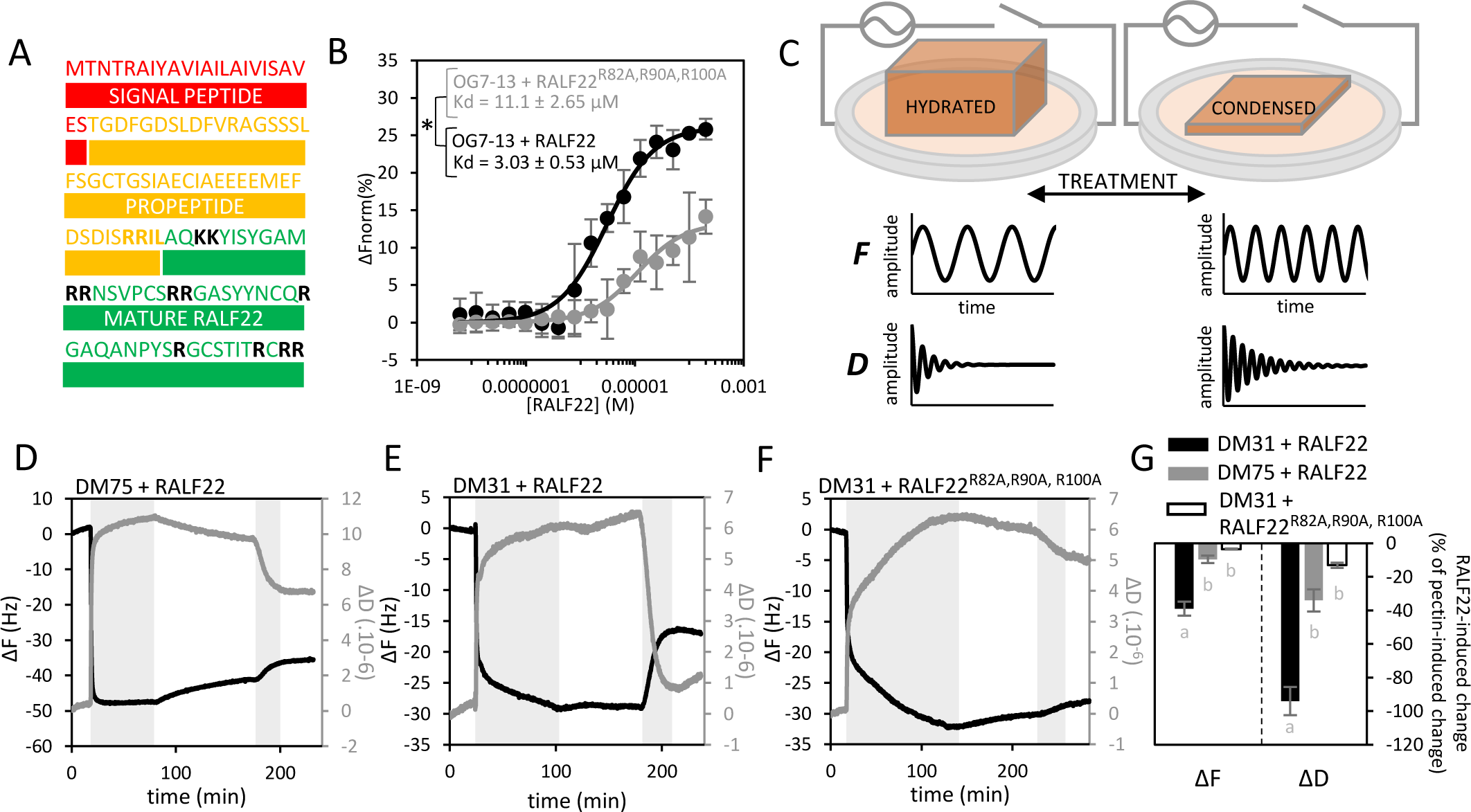
RALF22 binds and compacts demethylesterified homogalacturonan. (A) mature RALF22 is a positively charged protein. In bold: the amino acids which are positively charged at pH 7.0. (B) MST affinity plots showing that RALF22 interacts with OG7-13, a 7-13mer long fully demethylesterified oligogalacturonide (n=3). Mutating three cationic arginines on the RALF22 (RALF22^R82A,R90A, R100A^) surface leads to a 3.7-fold reduction in affinity (n=3). (C) illustration of the QCM-D working principle. An alternating current is applied to a quartz surface which holds the pectin gel. The frequency of the lateral quartz oscillation (F) is inversely correlated with the mass of the pectin layer. The time it takes for the lateral oscillation to dissipate (D) when the current is interrupted positively correlates with the rigidity of pectin layer. (D-F) Representative ΔF and ΔD QCM-D traces for a pectin layer with a degree of methylesterification of 75% (D) or 31% (E) treated with 1µg.mL^-1^ RALF22 or DM31 pectin treated with 1µg.mL^-1^ RALF22^R82A,R90A, R100A^. Grey zones highlight the intervals during which pectin or RALF were applied (cfr. Fig. S4A,B). (G) Quantification of the % change in ΔF and ΔD upon RALF treatment for the conditions presented in D-F (DM75 + RALF22, n=14; DM31 + RALF22, n=14; DM31 + RALF22^R82A,R90A, R100A^, n=2). Data is shown as the mean ± SEM. Different letters represent statistical significance (α=0.05).

### RALF22 forms periodic circumferential rings in the root hair cell wall

To investigate how endogenous RALF22 regulates cell expansion, we generated *ralf22-2* lines expressing mCherry-tagged mature RALF22 (*ralf22-2 x pRALF22::mCherry-RALF22_mature_*; see Star Methods). *Ralf22-2* x *pRALF22::mCherry-RALF22_mature_* seedlings had RHs that were indistinguishable from Col-0, showing that the tagged protein is fully functional (Fig. 4A, B). Surprisingly, mCherry-RALF22 accumulated in the RH CW, where it formed regularly spaced microdomains (Fig. 4C), which remained immobile throughout RH development (Fig. 4C, D, movie 3A). These microdomains arise at the RH tip, the exclusive site of RALF22 secretion (Fig. 4E, movie 3B). Indeed, using Fluorescence Recovery After Photobleaching (FRAP) we found that mCherry-RALF22 fluorescence rapidly recovered at the tip, but did not recover subapically (Fig. 4F, movie 4). At the growing apex, *de novo* mCherry-RALF22 deposition oscillated in antiphase with the growth rate oscillations, illustrating that temporal growth rate periodicity is translated into spatial periodicity in RALF22 microdomain formation in the CW (Fig. 4G). Closer inspection of the mCherry-RALF22 microdomains, using spinning disk microscopy, revealed that they represent regularly spaced rings surrounding the RH’s tubular structure (Fig. 4H). These rings first appear at the tip, after which they circumferentially expand 11.5±0.9-fold (n=8) in the expanding dome.

**Figure 4.**
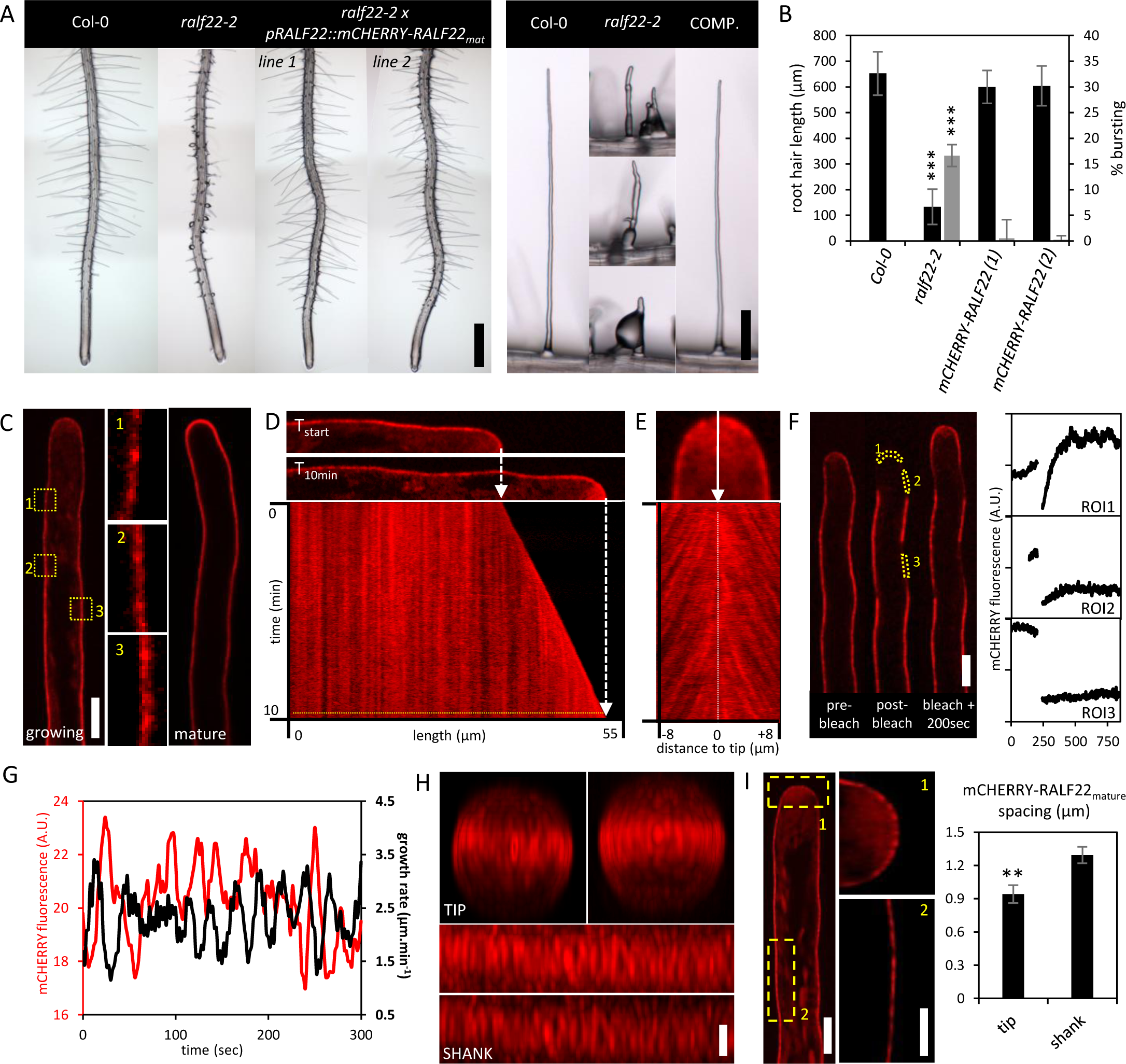
RALF22 forms circumferential rings in the root hair cell wall which illustrate anisotropic wall expansion. (A) representative images of 6-day-old Col-0, *ralf22-2* and *ralf22-2 x pRALF22::mCHERRY-RALF22_mature_* roots (scale bar=500µm) and RHs (scale bar=100µm). (B) quantification of the average RH length and percentage of RH bursting for each genotype (n≥5, ***p<0.001). (C) Representative longitudinal optical section of a growing (left) and a mature (right) *ralf22-2 x pRALF22::mCHERRY-RALF22_mature_* RH (scale bar=8µm). ROIs showing the presence of mCHERRY-RALF22_mature_ microdomains in the RH CW (scale bar=2µm). (D) maximal projection kymograph corresponding to a 10min timelapse acquisition of a growing *ralf22-2 x pRALF22::mCHERRY-RALF22_mature_* RH. Vertical red lines represent mCHERRY-RALF22_mature_ microdomains that remained stationary in the CW throughout the acquisition. The longitudinal fluorescence pattern for quantification of the mCHERRY-RALF22_mature_ microdomain spacing was extracted along the yellow dotted line. (E) maximal projection kymograph of the RH tip CW after removal of the dome’s concave shape. Red lines represent stationary mCHERRY-RALF22_mature_ microdomains, which arise at the very apex and move towards the shank as the tip grows forward. (F) Representative snapshots of a growing *ralf22-2 x pRALF22::mCHERRY-RALF22_mature_* RH before, upon and after photobleaching of mCHERRY-RALF22_mature_ in ROIs in the CW of the expanding dome, below the tip and in the shank (scale bar=5µm). Graphs showing mCHERRY fluorescence recovery at the apex only. (G) representative timeseries showing that RH growth rate and apical mCHERRY-RALF22_mature_ fluorescence oscillate in antiphase. (H) frontal and lateral high-resolution z-projections of mCHERRY-RALF22_mature_ organization in the RH CW, illustrating the formation of regularly spaced mCHERRY-RALF22_mature_ rings (scale bar=2µm). (I) Representative high resolution longitudinal optical section of a representative growing RH (scale bar=5µm) showing mCHERRY-RALF22_mature_ microdomains in the tip and the shank (scale bar=4µm). Comparative analysis of mCHERRY-RALF22_mature_ microdomain spacing in the tip (n=12) vs. the shank (n=15). Data is presented as the mean ± SEM. Asterisks depict statistical significance (**p<0.01).

To better understand this localization pattern, we compared the spatial periodicity of mCherry-RALF22 rings along the RH’s longitudinal axis with the temporal periodicity in RH growth rate (Fig. S5A-D). Based on wavelet transformation of the longitudinal fluorescence pattern present in the mCherry-RALF22 kymographs (e.g. Fig. 4D) the average RALF22 ring spacing was 1.34±0.08 µm (Fig. S5A, B). Interestingly, for the same RHs, a single growth rate period (30.47±0.93s at 2.15±0.06 µm.min^-1^) attributed to an average RH length gain which was 1.21-fold lower (1.09±0.04 µm.period^-1^; Fig. S4C-E), suggesting that the spacing between RALF22 rings increases when moving from tip to shank. We then used wavelet analysis to compare the mCherry-RALF22 ring spacing in the expanding dome and along the RH shank (Fig. 4I, Fig. S5F-K). Indeed, whilst the spacing in the tip was 0.94±0.08 µm, shank-localized rings were spaced 1.37-fold further apart at 1.29±0.08 µm (Fig. 4I, Fig. S5J, K).

Together, these data show that, during RH growth, the temporal periodicity in growth rate translates into the spatial periodicity in RALF22 ring formation in the CW. In addition, wall expansion over the surface of the dome was highly anisotropic as shown by the 8.4x larger circumferential strain (11.5-fold increase) of the rings relative to the meridional (from tip to shank) strain (1.37-fold increase).

### RALF22 forms a complex with LRX1 in root hair cell walls

The periodic arrangement of RALF22 in the RH CW suggests that it could associate with other CW epitopes with a similar ring-like organization. Interestingly, RALF-mediated regulation of CW integrity depends on two distinct receptor families, the biological relevance of which is unclear. Hence, RALF peptides have previously been shown to bind CW proteins of the LEUCINE-RICH REPEAT EXTENSIN (LRX) family, with distinct and mutually exclusive binding modes to the LLG-CrRLK1L signaling system^23,27,29–31^. In RHs, LRX1 and LRX2 have partially redundant roles in sustaining RH growth^51^. The RH phenotype of the *lrx1-1/2-1* loss-of-function mutant is indistinguishable from *ralf22*, suggesting that they could be functionally related (Fig. 5A, B, movie 5). To investigate the plausibility of a RALF22-LRX1/2 complex, we generated a peptide docking model using the crystal structure of the LRR domain of LRX2 (LRX2, PDB: 6QXP)^29^ and an AlphaFold2 model of the corresponding LRX1_LRR_ domain (Fig. 5C). RALF22_mature_ was modelled using the crystal structure of RALF4 (PDB: 6TME) as a reference. Similar to the LRX8_LRR_-RALF4 complex^29^, we found that the conserved binding pockets of LRX2 and LRX1 could accommodate RALF22_mature_ (Fig. 5C, LRX2_LRR_-RALF22; total energy score=-66.43 kcal.mol^-1^; root mean square deviation=0.760 Å). LRX1_LRR_-RALF22 and LRX2_LRR_-RALF22 dimeric complexes could indeed be purified from insect cells that co-expressed LRX1_LRR_ or LRX2_LRR_ with RALF22 (Fig. 5D). These complexes were extremely stable as shown by the inability of LRX1_LRR_-RALF22 to dissociate at pH 2.0 (Fig. 5D, E) and their 92.3°C (LRX1_LRR_-RALF22) and 83.8°C (LRX2_LRR_-RALF22) dissociation temperatures in thermal shift assays (Fig. 5F). These results suggest that the LRX1_LRR_-RALF22 and LRX2_LRR_-RALF22 are bona fide complexes with even higher stability than the previously described LRX8_LRR_-RALF4 pollen expressed complex^29^.

**Figure 5.**
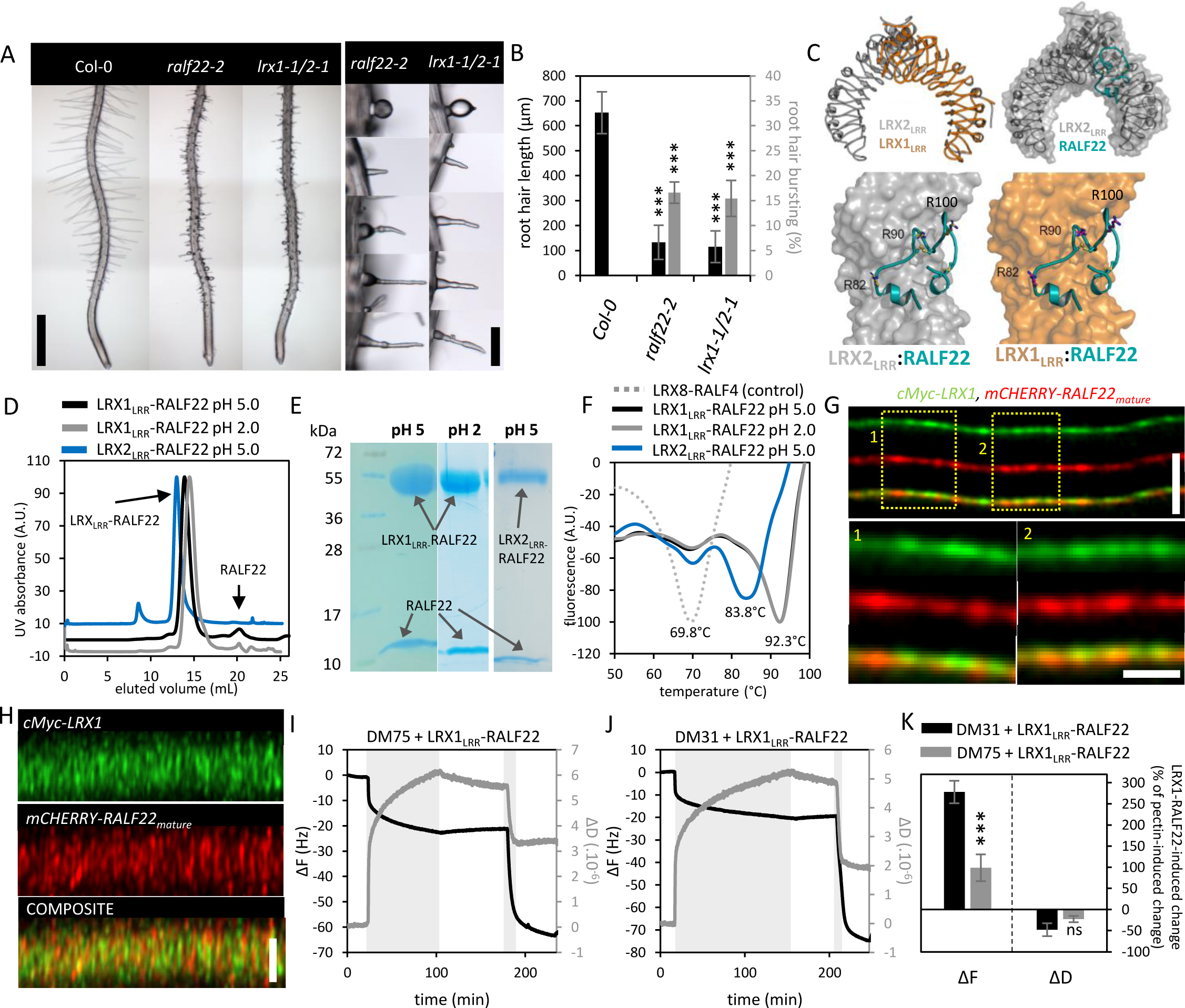
RALF22 binds to LRX1 in the root hair cell wall and compacts demethylesterified homogalacturonan. (A) The *lrx1-1/2-1* and *ralf22-2* phenotypes are indistinguishable. Representative pictures (scale bar=500µm) and close-ups (scale bar=100µm) of 6-day-old seedlings are shown. (B) quantification of the RH length and % of RH bursting of Col-0, *ralf22-2* and *lrx1-1/2-1* seedlings (n≥5, ***p<0.001). (C) Structural superimposition of the LRX2 (PDB:6QXP; grey) and LRX1 AlphaFold2 model (orange) (top left). Root Mean Square Deviation (R.M.S.D) of 0.15Å comparing 342 pairs of corresponding Cα atoms between the two proteins. Full molecular docking model of RALF22 (green) with LRX2_LRR_ (PDB: 6QXP; grey) (top right). Enlarged view of the peptide docking models showing RALF22 (green) in the LRX2_LRR_ and LRX1_LRR_ (AlphaFold2 model) binding pockets (bottom). The three exposed Arginines (R82, R90, R100) required for pectin binding are depicted as sticks. (D) Size exclusion chromatography of LRX1_LRR_-RALF22 (pH 5.0 and 2.0) and LRX2_LRR_-RALF22 (pH 5.0) purified from insect cells, showing that the complex does not dissociate at pH 2. (E) SDS protein gels of the fractions corresponding to the SEC elution peaks for LRX1_LRR_-RALF22 (pH 5.0 and 2.0) and LRX2_LRR_-RALF22 (pH 5.0). (F) thermoshift assay of LRX8_LRR_-RALF4 (control), LRX1_LRR_-RALF22 and LRX2_LRR_-RALF22. LRX1_LRR_-RALF22 was subjected to pH 5.0 and pH 2.0. Low pH does not affect the thermostability of the complex. (G) representative longitudinal optical sections of the *lrx1-1 x pLRX1::cMyc-LRX1 x pRALF22::mCHERRY-RALF22_mature_* CW in which LRX1 was labeled with a green fluorescent anti-cMyc antibody (scale bar=5µm). Dots represent cMyc-LRX1 (green) and mCHERRY-RALF22_mature_ microdomains (red). Composite images and magnifications (scale bar=1µm) show colocalization (yellow) between LRX1 and RALF22. (H) Representative lateral z-projections of cMyc-LRX1 (green) and mCHERRY-RALF22_mature_ (red) organization and colocalization (yellow) in the RH CW (scale bar=5µm). cMyc-LRX1 colocalizes with RALF22_mature_ rings in the CW. (I,J) representative ΔF and ΔD traces of DM75 (I) and DM31 (J) treated with 10 µg.mL^-1^ of LRX1 -RALF22. (K) Quantification of the % change in ΔF and ΔD (relative to the pectin-induced change) upon LRX1_LRR_-RALF22 treatment for the conditions presented in I and J (DM31 + LRX1_LRR_-RALF22, n=11; DM75 + LRX1_LRR_-RALF22, n=6). Data is shown as the mean ± SEM. Asterisks represent statistical significance (p-value; ***<0.001).

To study the LRX1-RALF22 interaction *in planta*, we introduced *pRALF22::mCHERRY-RALF22_mature_*into *lrx1-1* plants that express a functional cMyc-tagged LRX1 (*lrx1-1 x pLRX1*::*cMyc-LRX1*^52^). cMyc-LRX1 also formed rings perpendicular to the RH’s longitudinal axis, that colocalized to a great extent with mCHERRY-RALF22_mature_ (Fig. 5G, H; Costes p-value>0.95; thresholded Manders’ coefficient (tM)_mCHERRY-RALF22[lcMyc-LRX1_=0.71±0.22; tM_cMyc-LRX1[lmCHERRY-RALF22_=0.70±0.21; n=11). These data are consistent with the presence of LRX1-RALF22 complexes *in planta* but are expected to be incomplete given the presence of endogenous RALF22 and LRX2. Together, these data show that RALF22 adopts a periodic pattern in the RH CW at least in part as an LRX1-RALF22 complex.

### LRX1-RALF22 also binds and compacts demethylesterified homogalacturonan

Next, we investigated whether pectin binding is preserved in the LRX1_LRR_-RALF22 complex. A possible pectin interaction is suggested by the 3D crystal structure of the LRX8_LRR_-RALF4 dimer, in which RALF4 displays a highly positively-charged and surface-exposed patch of amino acids^29^. Homology modeling shows that this is also true for the LRX1_LRR_-RALF22 complex, albeit with a less pronounced surface charge compared to LRX8_LRR_-RALF4 (Fig. 5C), comprising the three arginines (R82, R90, R100) that are critical for free RALF22-pectin binding (Fig. 3B, F, G, Fig. 5C). QCM-D indeed showed an interaction of the LRX1_LRR_-RALF22 complex with DM75 and DM31 pectin layers (Fig. 5I-K). In contrast to free RALF22 however, we observed a charge-dependent mass increase (decrease in ΔF_LRX1-RALF22_ for DM31 of 278.21±26.64%, n=11; decrease in ΔF_LRX1-RALF22_ for DM75 of 98.62±31.52%, n=6) (Fig. 5I-K). Together with the mass increase, we observed a decrease of the viscoelasticity (ΔD_LRX1-RALF22_ decrease for DM31 of 48.02±15.06%, n=11; ΔD_LRX1-RALF22_ decrease for DM75 of 22.54±7.37%, n=6), indicating that LRX1_LRR_-RALF22, like free RALF22, promoted the stiffening of the pectin layers (Fig. 5K). Concerning the LRX1_LRR_-RALF22-dependent mass increase, as opposed to the mass decrease observed upon free RALF22 supply, this is most likely due to the much higher mass of the LRX1_LRR_-RALF22 dimer (94.18kDa) compared to the mass of RALF22 alone (5.6kDa), which most likely exceeds the mass loss due to pectin dewatering. The observed decrease in ΔD is presumably the resultant of the stiffening effect of the LRX1_LRR_-RALF22-pectin interaction and the decrease in viscoelasticity due to the addition of extra mass to the layer. More detailed studies are needed to quantify these different effects. Nevertheless, these results show that LRX1_LRR_-RALF22, like free RALF22, is able to bind and stiffen pectin in a charge-dependent manner.

### RALF22 is required for the self-assembly of pectic homogalacturonan

Given the charge-dependence of the interaction of RALF22 and LRX1-RALF22 with pectin, pectin demethylesterification is expected to be critical for the assembly and expansion of the RH CW. To investigate this, we first supplemented growing RHs with the PME inhibitor EGCG, while imaging [Ca^2+^]_cyt_, pH_ext_ and CW charge (Fig. S6; movie 6). EGCG treatment induced a short decrease in growth rate, followed by a transient growth spurt and a transition from longitudinal to radial expansion (=swelling) at the tip (Fig. S6A; movie 6). This response was accompanied by a transient increase in propidium iodide staining, a dye that stains charged polymers in the CW^53^, and high amplitude [Ca^2+^]_cyt_ and pH_ext_ transients that coincided with rapid tip swelling (Fig. S6A; movie 6). No such changes were observed upon mock treatment (Fig. S6B, movie 6). Together these observations underscore the dynamic nature of pectin metabolism in growing RHs, where rapid feed-back signaling compensates the effect of the inhibition of PME activity, at least in part through the alkalinization of the cell surface. The latter is expected to promote PME activity, given the high pH optimum of plant PMEs^34^. The radial expansion of the RH tip as rapid as one minute after EGCG treatment emphasizes the critical role for pectin demethylesterification in maintaining polar growth of RHs.

Next, we hypothesized that preventing pectin demethylesterification in the RH CW would also affect the cell’s response to RALF22 treatment. To test this, we measured [Ca^2+^]_cyt_ responses upon RALF22 treatment, in the presence or absence of EGCG, in seedlings expressing the G5A BRET sensor driven by the RH-specific promoter pEXPa7^54^. This assay, in contrast to the observation of individual RHs in microfluidic chips (Figure 2; movie 2), is more suitable for gathering quantitative data from large numbers of treatments. We found that a treatment as short as 30 min, at an EGCG concentration close to the EC_50_ for root growth inhibition (32 µM^55^), greatly reduced the subsequent RALF22-induced [Ca^2+^]_cyt_ transient (Fig. S6C, D), showing that, also in vivo, pectin demethylesterification is crucial for RALF22 function.

To further investigate the *in vivo* connection between the methylesterification status of HG and the periodic organization of (LRX1-)RALF22 in the CW, we labeled *ralf22-2* x *pRALF22::mCherry-RALF22_mature_* RHs with monoclonal antibodies (mABs) directed against different HG epitopes (Fig. 6, Fig. S7). LM20, which labels methylesterified HG^56^, was enriched at the tip and labeled sparse patches along the RH shank (Fig. S7A-C), corroborating that methylesterified HG is deposited and demethylesterified in the expanding dome. Interestingly, LM20 labeled an outer layer of the CW that was devoid of mCHERRY-RALF22_mature_, which localized closer to the plasma membrane both in the tip and the shank (Fig. S7A, B). The mAb 2F4, an IgG1 that binds to Ca^2+^-crosslinked demethylesterified HG^57^, labeled the RH tip and shank and, like LM20, preferentially labeled an outer layer of the CW, which only partially overlapped with mCherry-RALF22_mature_ (Fig. S7D, E; Costes p-value>0.95; tM_mCHERRY-RALF22[l2F4_=0.56±0.04; tM_2F4[lmCHERRY-RALF22_=0.49±0.02; n=8). Also along the shank, Ca^2+^-crosslinked demethylesterified HG was organized in rings, where regions with a relatively lower mCherry-RALF22_mature_ abundance appeared to show higher 2F4 labeling and vice versa (Fig. S7E, F). Finally, PAM1, a small HIS-tagged scFv mAb directed towards long (>30DP) stretches of block-wise demethylesterified HG^58^, labeled the RH CW both in the tip and shank (Fig. 6A-D). In contrast to LM20 and 2F4 epitopes, this epitope revealed circumferential rings in the RH CW that largely overlapped with mCherry-RALF22_mature_ (Fig. 6A-D; Costes p-value>0.95; tM_mCherry-RALF22 PAM1_=0.74±0.05; tM_PAM1 mCherry-RALF22_=0.74±0.04; n=12). These results, in accordance with the charge-dependence of the (LRX1-)RALF22-pectin interaction (Fig. 3B, D-G, Fig. 5I-K, Fig. S6C, D), show that newly secreted pectin, upon demethylesterification, interacts with (LRX1-)RALF22 *in muro* where it becomes part of periodic rings (Fig. 6A-D).

**Figure 6.**
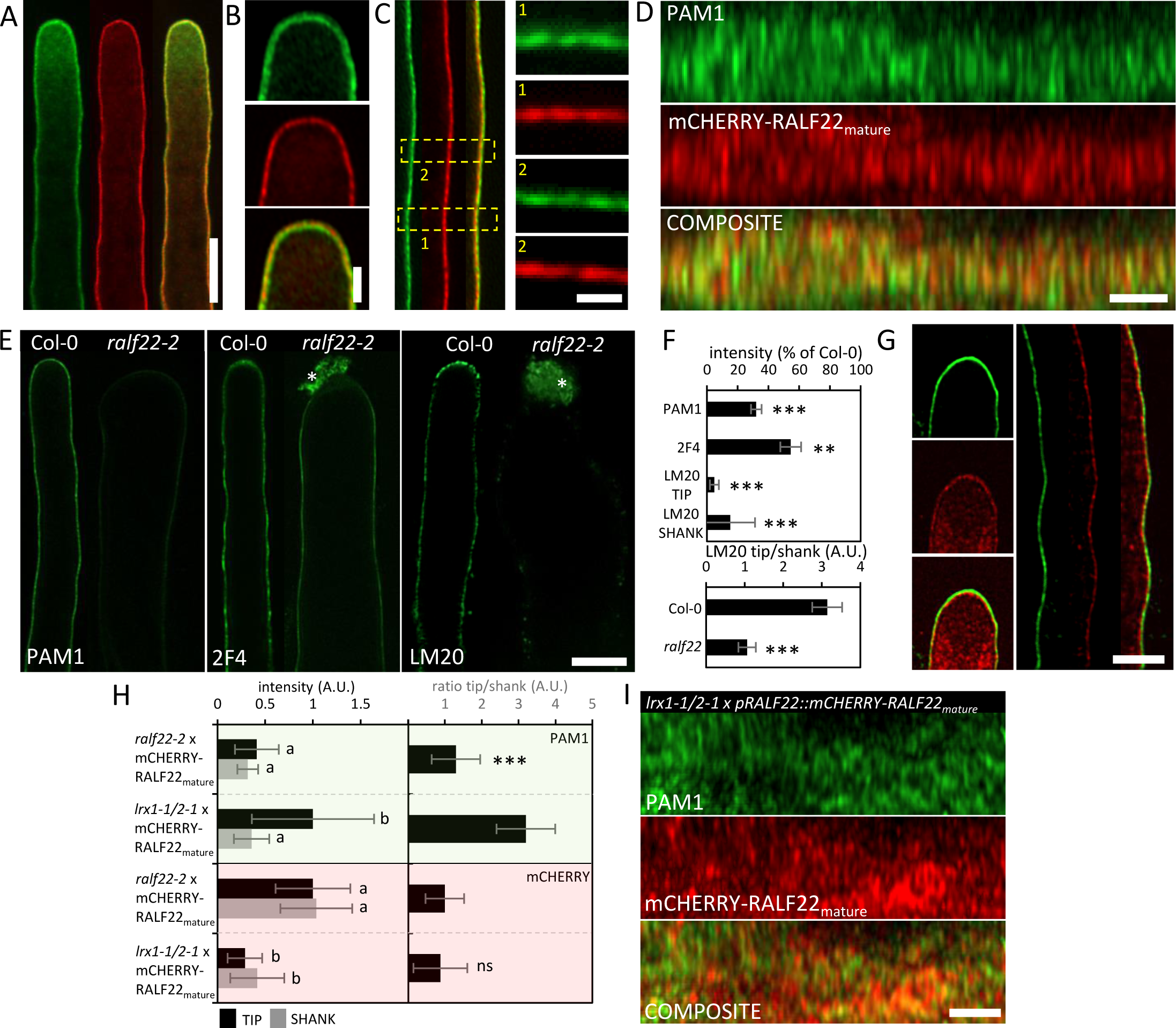
(LRX1-)RALF22-homogalacturonan interaction regulates the organization of the pectic root hair cell wall. (A-D) representative images showing the colocalization of PAM1 (>30DP long stretches of block-wise demethylesterified homogalacturonan) and mCHERRY-RALF22_mature_ in the RH tip and shank. (A) longitudinal optical sections of a PAM1 (green) and mCHERRY-RALF22_mature_ labeled RH (scale bar=10µm), which was fixed while growing. Yellow indicates colocalization in the composite image. (B) close-ups of RH tip presented in A showing that PAM1 and mCHERRY-RALF22_mature_ colocalize along the growing dome (scale bar=2.5µm). (C) representative longitudinal optical sections of the shank CW and ROIs showing the presence of overlapping microdomains of demethylesterified HG (PAM1; green) and mCHERRY-RALF22_mature_ (red) (scale bar=1µm). (D) lateral z-projections of the RH shank CW showing the formation of colocalized PAM1 and mCHERRY-RALF22_mature_ circumferential rings (scale bar=5µm). (E) representative longitudinal optical sections of ralf22-2 RHs labeled with PAM1 (demethylesterified HG), 2F4 (Ca^2+^-crosslinked demethylesterified HG) and LM20 (methylesterified HG) (scale bar=10µm). Asterisks indicate expelled intracellular debris as a result of cell bursting. (F) quantification of the labeling intensities for the PAM1 (n_col-0_=12; n_PAM1_=16), 2F4 (n_col-0_=8; n_2F4_=9) and LM20 epitopes (n_col-0_=19; n_PAM1_=9) relative to Col-0 (top barplot), and the ratio between between tip and shank methylesterified HG abundance (LM20; lower barplot). Data is shown as the mean ± SEM. Asterisks represent statistical significance (p-value; **<0.01; ***<0.001). (G) representative longitudinal optical sections of the tip and shank CW of a surviving PAM1 (green)-labelled lrx1-1/2-1 RH expressing mCHERRY-RALF22_mature_ (red). Yellow indicates colocalization in the composite frame (scale bar=5µm). (H) Quantification of the PAM1 (green) and mCHERRY-RALF22_mature_ (red) CW labeling intensities in the tip and shank of *lrx1-1/2-1 x pRALF22::mCHERRY-RALF22_mature_* and complemented *ralf22-2 x pRALF22::mCHERRY-RALF22_mature_* RHs (left half) alongside the tip/shank labeling ratio (right half). Data is shown as the mean ± SEM. Different letters and asterisks represent statistical significance (p-value; ***<0.001). (I) lateral z-projections of the PAM1 labeled *lrx1-1/2-1 x pRALF22::mCHERRY-RALF22_mature_* RH shank CW showing the fragmented pattern of demethylesterified HG and RALF22 organization (scale bar=5µm).

The periodic CW pattern could indicate that mCherry-RALF22_mature_ associates with a preexisting HG pattern in the CW or that the (LRX1-)RALF22 interaction with demethylesterified HG could be responsible for inducing the formation of periodic pectin rings. To investigate this possibility, we labeled the CW of *ralf22-2* RHs with anti-HG antibodies (Fig. 6E, F). Interestingly, in *ralf22-2,* the abundance of block-wise demethylesterified HG (PAM1) and Ca^2+^-crosslinked demethylesterified HG (2F4) was reduced by 68±3% and 46±7% respectively, relative to Col-0 (Fig. 6E, F). In addition, methylesterified HG (LM20), showed a 95±3% and 85±16% reduced labeling in the *ralf22-2* tip and shank, respectively, as compared to Col-0 (Fig. 6E, F). Combined, these data illustrate that, in the absence of RALF22, overall pectin secretion is reduced and/or turnover is enhanced.

We then investigated the requirement of LRX1/2 for the periodic organization of the pectic CW by studying the RHs of the double *lrx1-1/2-1* mutant expressing mCherry-RALF22_mature_ (*lrx1-1/2-1 x pRALF22::mCherry-RALF22_mature_*, Fig. 6G-I, movie 7). These RHs, like those of the untransformed *lrx1-1/2-1* parent, grew slowly and frequently burst. In surviving *lrx1-1/2-1* RHs, most mCherry-RALF22_mature_ accumulated in intracellular compartments, yet a small amount was still secreted to the CW (Fig. 6G, movie 7). CW labeling was 3.5- and 2.5-fold weaker in the tip and shank CW respectively, compared to the complemented *ralf22-2 x pRALF22::mCherry-RALF22_mature_* line (Fig. 6G, H, movie 7). Interestingly, in the absence of LRX1 and LRX2, mCherry-RALF22_mature_ and PAM1 organization showed a much more fragmented pattern compared to the control (Fig. 6D, I). Moreover, in the *lrx1-1/2-1* CW, demethylesterified HG accumulated 2.4-fold, relative to the WT, at the tip, but not in the shank (Fig. 6G, H). Given that pectin secretion occurs at the tip, this suggests an increased pectin secretion and turnover in the absence of LRX1/2. Moreover, the PAM1 epitope and mCherry-RALF22_mature_ showed only limited colocalization in the shank of *lrx1-1/2-1* RHs (Fig. 6I; Costes p-value>0.95 for 4 out of 6 samples; tM_mCherry-RALF22 PAM1_=0.25±0.08; tM_PAM1mCherry-RALF22_=0.19±0.07; n=6). Together, these results show that the LRX1/2-RALF22 complex, through its binding and compaction of demethylesterified pectin, organizes the patterning of the pectic RH CW. In the absence of this patterning, pectin secretion and turnover are increased and the CW’s integrity is compromised.

## Discussion

Here, we identified RALF22 as the main trichoblast-expressed RALF peptide controlling growth and CW integrity of RHs. RALF22 binds LRX1 and LRX2, forming extremely stable complexes (dissociation t° > 80°C) *in vitro* (Fig. 5D-F) and, given the colocalization pattern (Fig. 5G-H), most likely also *in vivo*. RALF22 also binds LLG1 and recruits FER into an active membrane-bound ternary signaling complex (Fig. 2B, C). The *in vivo* relevance of this interaction in RHs is shown by the requirement of FER to induce sustained surface alkalinization and growth arrest upon RALF22 treatment (Fig. 2E-J, movie 2).

What is the functional relevance of the existence of two distinct, mutually exclusive binding partners^29^ for the same peptide at the surface of the same cell? This report provides an answer to this key question by showing that RALF22 is not only a signaling peptide, but that it also has a structural role (Fig. 7). Indeed RALF22, as a free peptide or bound to LRX1, binds demethylesterified pectin *in vitro* (Fig. 3B-G, Fig. 5I-K). Pectin binding is also relevant *in vivo*, as suggested by (i) the diminished [Ca^2+^] response upon RALF22 treatment of RHs pretreated with EGCG (Fig. S6C, D), (ii) the colocalization of mCHERRY-RALF22 with blockwise demethylesterified HG (PAM1 epitope; Fig. 6A-D), but not with highly methylesterified HG (LM20 epitope; Fig. S7A-C), in the RH CW, where it forms stable microdomains organized in periodic concentric rings (Fig. 4C-I, Fig. S5) and (iii) the loss of proper pectin organization in the absence of RALF22 or LRX1/2 (Fig. 6E-I).

**Figure 7.**
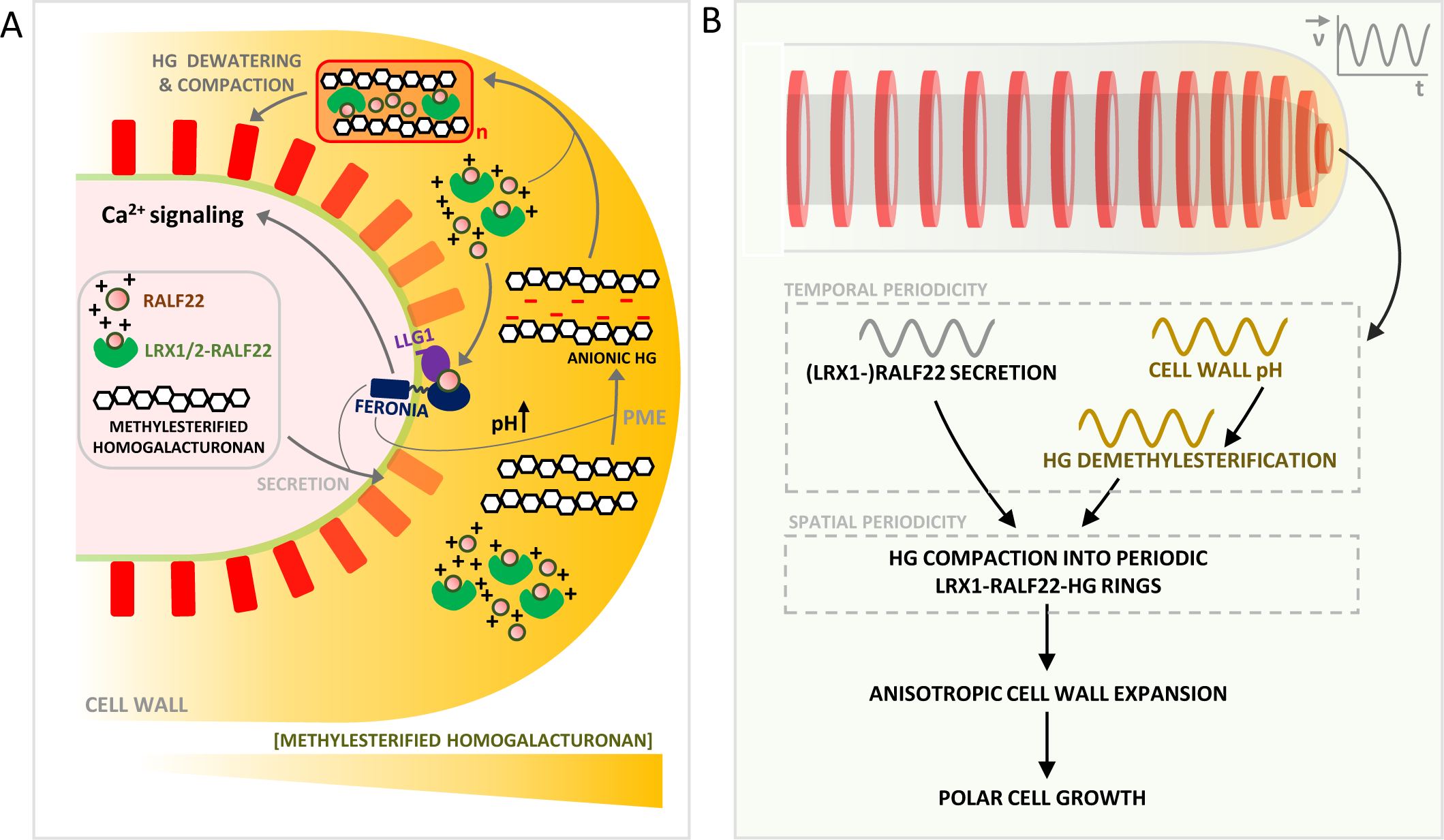
Model of the dual structural and signaling role of RALF22 in the periodic assembly of the root hair cell wall. (A) RALF22, LRX1-RALF22 and methylesterified HG are secreted at the growing RH tip. RALF22 forms a ternary complex with LLG1 and FER, thereby regulating its own secretion and inducing downstream cytosolic calcium signaling and alkalinization of the apical CW. In the CW, Pectin Methyl Esterases (PMEs), which have a high pH optimum, catalyze HG demethylesterification, generating anionic HG through the formation of stretches of negatively charged carboxyl groups. RALF22 and LRX1-RALF22, electrostatically interact with poly-anionic demethylesterified HG. This induces HG dewatering and compaction, and the self-assembly of the HG matrix into (LRX1-)RALF22-HG microdomains. (B) These microdomains represent periodic circumferential (LRX1-)RALF22-HG rings which originate in the growing tip, and arise as a consequence of oscillatory (LRX1-)RALF22 secretion at the tip and apical pH oscillations which catalyze periodic HG demethylesterification. This temporal periodicity is translated towards the spatial periodicity of (LRX1-)RALF22-HG ring assembly. (LRX1-)RALF22-HG rings illustrate highly anisotropic CW expansion, in which the circumferential strain of the rings greatly surpasses the meridional strain, which is a prerequisite for polar RH growth.

A mechanism for microdomain formation is suggested by the capacity of free RALF22 and LRX1_LRR_-RALF22 to induce the compaction of a pectin layer. The charge-dependence of this interaction suggests that the free unstructured and positively-charged peptide induces water loss through the neutralization of the pectin charges and the resulting loss of osmotically active counterions in the pectin gel. The interaction of the LRX1_LRR_-RALF22 hetero-tetramer with pectin may be more complex, since here the peptide is folded and exposes a structured surface with positively charged residues that, besides electrostatic interactions, may also recognize structural features of the polysaccharide, such as charge patterns, as reported for instance for GAG-binding proteins in the animal ECM^1,59^, or acetyl^60^ or xylosyl^61^ substitution motifs. In addition, the LRX1_LRR_-RALF22 dimer may promote polymer crosslinking, thus reducing the water binding capacity (Fig. 3D-G, Fig. 5I-K, Fig. S4A, B)^62^. Compaction of GAGs by GAG cross-linking proteins creates denser zones in the gel, but also augments the porosity of the non-compacted zones as observed during the formation of perineural nets, the porous aggregated matrices surrounding neurons^16^. A similar process may explain why RALF22 supplementation triggers the migration of dextran-coupled FITC and TRITC to the RH CW (Fig. 2E, F, J, movie 2). It should be noted that, besides the RALF22-binding LRR domain, LRX1 also comprises a C-terminal extensin domain. At least one extensin family member, AtEXT3, was able to polymerize into a scaffold *in vitro* thanks to its amphiphilic properties^63^. The authors hypothesized that this scaffold could form coacervates with pectin, due to its repeated basic domains, and thus structure the CW. This polymer network could then be consolidated by intermolecular oxidative crosslinking of the extensin’s tyrosine residues. Interestingly, since the extensin domain of LRX1 itself does not have such basic domains, RALF22 may play this pectin-recruiting role in the complex. The advantage of such a modularity might be twofold: first the same LRX could bind different CW polysaccharides or polysaccharide motifs by switching RALF isoforms, perhaps promoting different CW architectures, and second, free RALF peptide may be part of a FER-dependent feedback signaling mechanism that coordinates its own incorporation into the CW.

This study provided striking new insights into the logic of RALF feedback signaling. Whereas RALF22 treatment resulted in a dramatic change in the physicochemical properties of the CW, the cell required FER to induce a normal pH and growth response (Fig. 2E-J, Fig. S3B-E, movie 2). One way to explain these results is by assuming that the LLG1-RALF22-FER complex acts as a mechano-sensor, which responds to CW compaction induced by (LRX1-)RALF22. In other words, (LRX1-)RALF22 at the same time creates a signal (CW compaction) and controls the sensitivity of the detection system, possibly through LLG1-RALF22-FER recruitment. A role for LLG1/FER in mechano-sensing has been proposed previously^64–66^. For instance, stretched epidermal cells upon root bending displayed biphasic cytosolic Ca^2+^ responses, with a rapid transient and a slower more sustained component. Only the latter disappeared in *llg1* and *fer* mutants^65^, whereas the initial transient might come from another mechanosensor, such as a mechanosensitive Ca^2+^ -permeable channel^67^. Similarly in RHs, we observed that in *fer-4*, RALF22 still induced a transient growth inhibition and surface pH increase at the tip, after which growth, pH and [Ca^2+^] oscillations resumed, in contrast to the wild type, where the surface alkalinisation propagated along the shank and growth did not recover (Fig. 2E-J, Fig. S3B-E, movie 2). Mechanosensing in the leaf epidermis also appears to depend on FER, where it signals through the activation of ROP6 by FER-bound ROPGEF14^65^. This in turn is thought to affect microtubule and actin dynamics and secretion. Interestingly, we observed an over 90% reduction in methylesterified (LM20) pectin at the RH tip of *ralf22* and an over two-fold increase of the epitope in *lrx1/2* (Fig. 6H, I). This suggests that free RALF22, which is absent in *ralf22* and expected to be more abundant in the absence of LRX1/2, may promote the secretion of pectin at the tip.

RALF22-induced and FER-dependent sustained alkalinization is also expected to promote HG demethylesterification given the alkaline pH optimum of PMEs^34^. Together, free RALF22 might signal the need for more polyanionic pectin as a substrate for the CW-structuring LRX1-RALF22-pectin complexes. In this context it is of interest to note that the spacing of the CW microdomains in the RH tip matched the length increase during a growth/surface pH oscillation cycle (Fig. 4I, Fig. S5), and that apical RALF22 deposition oscillated in antiphase with the growth rate (Fig. 4G). This supports the idea that CW assembly is a cyclic process in RHs along the following scenario (Fig. 7): first, wall material, including methylesterified HG and LRX1-bound and free RALF22, is deposited at the tip of the growing RH. A concomitant or subsequent increase in CW pH through the binding of free RALF22 to LLG1/FER, activates processive pectin methylesterase (PME) activity, thus exposing stretches of negative charges on the polymer. This creates the substrate for the interaction with the LRX1/2-RALF22 dimer causing the dewatering of the pectin gel and the formation of a ring-like structure of higher density surrounding the RH tip. This structure is organized in such a way that it favors CW expansion in the circumferential direction. This deposition-demethylesterification-compaction-expansion process is repeated during each growth cycle. Given that cellulose^68^, xyloglucan^69^ and expansins^70^ are essential for RH growth, it will be interesting to see if expansin-mediated acid growth contributes to the preferentially circumferential expansion of the ring and whether this involves purely remodeling of the existing structure or the insertion of additional material. Further studies using microscopy at high spatio-temporal resolution^11^, should provide more insights into this fascinating CW assembly process in RHs and most likely in other cell types in plants.

## Acknowledgments

This work was funded by the Research Foundation Flanders (FWO; grants 1225120N and G013023N), the University of Antwerp (UA; BOF-KP; DOCPRO4) and LASERLAB-EUROPE (grant 654148) to SS and KV; the Agence Nationale pour la Recherche (ANR), project “HOMEOWALL” to HH; and the University of Lausanne, the European Research Council (ERC) grant agreement no. 716358 and the Swiss National Science Foundation grant 310030_204526 to JS. The authors would like to thank Jozef Mravec for sharing OG7-13, and Christoph Ringli, Alice Cheung and Simon Gilroy for kindly donating lrx1-1/2-1 and lrx1-1 x pLRX1::cmyc-LRX1, fer-4 x pFER::FER-GFP, and Col-0 x 35S::GCaMP3 seeds, respectively. Tou Cheu Xiong in thanked for providing the G5A Ca^2+^ sensor line. Our gratitude also goes to CPKelco (Denmark) for donating industrial low and high methoxyl pectin samples. The IJPB benefits from the support of Saclay Plant Sciences-SPS (ANR-17-EUR-0007). This work has benefited from the support of IJPB’s Plant Observatory technological platforms. Access to fluorescence microscopy infrastructure was provided by the Antwerp Centre for Advanced Microscopy (ACAM, UA). The purchase of the PerkinElmer UltraVIEW Vox and Leica SP8 confocal microscopes was supported by FWO-HERCULES infrastructure grants; AUHA-09-001, AUHA-15-12. The Nikon Sora microscope was purchased with the support of an FWO mid-size infrastructure (I003420N) and FWO IRI grant (I000123N).

## Author contributions

S.S., K.V., H.H. and J.S. conceived the project. S.M. aided in conceptualizing the research based on his work on LRX8-RALF4. J.K.L. and J.S. designed, produced and characterized all recombinant proteins. C.B. provided technical assistance for protein production. E.F. and M.G. designed and performed calcium assays using the G5A BRET sensor. S.S. generated all crosses and performed the cloning of the pEXPa7::G5A sensor and RALF22-related constructs. S.S. optimized microfluidics for imaging and performed the phenotyping, staining and imaging of live cell and fixed-tissue samples. S.S. performed image analysis, bio-informatics and statistics. N.C. assisted with live cell imaging, cloning and sample preparations. D.B. assisted with cloning, phenotyping and sample preparation. S.S. and M.G. performed Microscale Thermophoresis analysis. H.H. and B.C. conceived the QCM-D analysis. T.L. and H.H. performed the QCM-D analysis. T.L. characterized the pectin solutions. E.B. and C.M. provided technical and conceptual assistance with the in vitro QCM-D and pectin work. H.A. aided in conceptualizing the work and provided assistance with sample preparation for phenotyping. A.B., A.C., D.S.C.D and J.F. aided in the visualization and analysis of oscillatory parameters in live cell imaging. J.S. generated 3D protein structures and performed peptide docking modelling. S.S., H.H. and K.V. wrote the manuscript.

## Declaration of interests

The authors declare no competing interests.

## Methods

### Plant materials

*Arabidopsis thaliana* ecotype Columbia-0 (Col-0; N1092), *ralf22-2* (GK_293H09; N428125; At3g05490), *fer-4* (GK_106A06; N69044; At3g51550)*, eru* (SALK_083442; N583442; At5g61350), and *llg1-2* (SALK_086036; N586036; At5g56170) seeds were obtained from the Eurasian Arabidopsis Stock Centre (uNASC). Genotyping of *ralf22-2* plants was performed by PCR using T-DNA (GTAGATTTCCCGGACATGAAGCCA) and gene-specific primers (FW: ACCGGTCAACCAGTTTCTGCAT, REV: TTCAACGCCTGCACCTAGTGAT). The *ralf22-1* line was generated using CRISPR-cas9. *Lrx1-1/2-1* (At1g12040, At1g62440) and *lrx1-1 x pLRX1::cmyc-LRX1* seeds were kindly donated by Prof. Christoph Ringli. *Fer-4 x pFER::FER-GFP* was kindly donated by Alice Chueng. Col-0 x 35S::GCaMP3 seeds were kindly provided by Prof. Simon Gilroy. *ralf22-1*, Col-0 x *pEXPa7::G5A*, *ralf22-2 x pRALF22::mCHERRY-RALF22_mature_, lrx1-1/2-1 x pRALF22::mCHERRY-RALF22_mature_* and *lrx1-1* x *pLRX1::cmyc-LRX1/pRALF22::mCHERRY-RALF22_mature_* lines were generated during this study. *Eru x pERU::ERU-GFP* was generated previously^42^.

### Growth conditions

Seeds were surface sterilized and sown on RH growth medium (3mM KNO_3_, 2mM Ca(NO_3_)_2_.4H_2_O, 0.5mM MgSO_4_.7H_2_O, 1mM NH_4_H_2_PO_4_, 1mg.mL-1 thiamine, 0.5 mg.mL-1 pyridoxine-HCl, 0.5mg.mL-1 nicotinic acid, 0.56mM myo-inositol, 25mM KCl, 17.5mM H_3_BO_3_, 1mM MnSO_4_.H_2_O, 1mM ZnSO_4_.7H_2_O, 0.25mM CuSO_4_.5H_2_O, 0.25mM (NH_4_)6MoO_24_.4H_2_O, 25mM Fe-Na EDTA, 0.8% gelrite or phytagel, 1% sucrose, 2.3mM MES at pH 5.7) for phenotyping or basal Murashige and Skoog (MS) medium (1% sucrose, 1% plant agar, 0.5g.L^-1^ MES at pH 5.8) prior to microfluidic preparation. For live cell imaging and immunolocalization assays, plants were transferred to microfluidic chips and grown overnight in liquid medium (0.1mM KCl, 0.1mM CaCl, 1mM NaCl, 1% sucrose, 0.5g.L^-1^ MES at pH 6.0). Imaging of extracellular pH oscillations was performed in unbuffered liquid medium at pH 6.0.

Seeds were stratified for 2-3 days in the dark at 4°C and plates or microfluidics chips were placed vertically in a growth chamber with standard growth conditions (16h light-8h dark, 22°C).

### Identification of RH-specific RALFs

Using the BAR ePlant Browser^71^, Root-specific transcription was assessed for all *RALFs* that are represented on Affymetrix ATH1 arrays. Root-expressed, trichoblast-specific *RALFs* were identified from single cell RNAseq data based on expression in cluster 15 (trichoblasts) in T-SNE plots^36^. For *RALF22*, a trichoblast pseudotime expression profile was generated, showing *RALF22* expression in function of the trichoblast’s distance to the meristem^37^.

### Immunolocalization

Four day old Col-0, *ralf22-2*, *ralf22-2 x pRALF22::mCHERRY-RALF22_mature_, lrx1-1 x pLRX1::cmyc-LRX1/pRALF22::mCHERRY-RALF22_mature_* and *lrx1-1/2-1 x pRALF22::mCHERRY-RALF22_mature_* seedlings were grown overnight in microfluidics chips^72^ and fixed for 10min in buffered liquid medium containing 10% acetic acid and 3.7% formaldehyde. All subsequent steps were performed in buffered liquid medium. Each channel was washed (3x5min) and incubated for 15min with 50mM NH_4_Cl to neutralize residual formaldehyde. After washing (3x5min), aspecific binding sites were blocked for 30min in 3% BSA and incubated overnight at 4°C in the presence of 3% BSA and a 20-fold (2F4, LM20) or 40-fold (PAM1) primary antibody dilution. Seedlings were washed (3% BSA, 3x5min) and incubated for 30min at RT with a 50-fold secondary antibody dilution (2F4; goat anti-mouse IgG-FITC, LM20; goat anti-rat IgM-Alexa^488^, PAM1; Rabbit anti-His-DyLight^488^, cMyc-LRX1; anti-c-Myc-Alexa^488^). Following a final washing step (3x5min), the chips with labeled seedlings were used for imaging on a Leica TCS SP8 confocal laser scanning microscope (Leica-microsystems).

### Microscopy

Six-day-old plants grown on RH growth medium were used for phenotyping. A Nikon AZ100 multizoom macroscope or Zeiss AxioZoom v16 were used to collect 2h long timelapse acquisitions or images of whole roots and individual RHs.

For fluorescence microscopy plants were grown overnight in uncoated IBIDI µ-slides VI 0.4 filled with liquid medium^72^. Each channel contained a single plant.

Col-0 x *pRALF22::GFP* plants were counterstained by injecting 10µM propidium iodide (PI) (in liquid medium) in each channel. Roots were imaged in xyz-mode using a Leica TCS SP8 confocal laser scanning microscope (Leica-microsystems) equipped with a white light laser, 2 HyD detectors and a 5x HC PL FLUOTAR dry objective (NA 0.15). GFP (excitation: 488nm, emission: 501-536nm and PI (excitation: 556, emission: 590-710nm) fluorescence were captured in bidirectional line-scanning mode at 8-bit. The z-spacing was set to 1µm-2µm.

The same system with a 63x HC PL APO CS2 water immersion objective (NA 1.20) was used to image [Ca^2+^]_cyt_, pH_ext_, growth rate and CW porosity or CW charge dynamics upon treatment with RALF22 or EGCG respectively. Col-0 and *fer-4* plants were grown in microfluidics chips in 220µL of unbuffered liquid medium. Prior to imaging, 160µL of medium was removed from each channel. 60µL of medium containing 0.29 mg.mL^-1^ FITC-110kDa dextran and 0.25 mg.mL^-1^ TRITC-20kDa dextran (for RALF22 treatment) or 0.29 mg.mL^-1^ FITC-110kDa dextran and 10µM PI (for EGCG treatment) was injected back into the channel. Plants were allowed to recover for 30min and were subsequently imaged for 10-20min at 30 frames.min^-1^. FITC/GCaMP3 (excitation: 488nm, emission: 501-548nm) and TRITC (excitation: 544nm, emission: 560-710nm) or PI (excitation: 556nm, emission: 590-710nm) fluorescence were captured using bidirectional line-by-line scanning in xyt-mode. After ∼4min of steady state growth channels were injected with 60µL of liquid medium containing either 5µM RALF22 or 100µM EGCG and imaged for another 6-16min.

Timelapse acquisitions of *fer-4 x pFER::FER-GFP, eru x pERU::ERU-GFP, ralf22-2 x pRALF22::mCHERRY-RALF22_mature_* and *lrx1-1/2-1 x pRALF22::mCHERRY-RALF22_mature_* were acquired in buffered liquid medium on an inverted Nikon Ti wide-field microsope equipped with a micro-lenses enhanced PerkinElmer UltraVIEW Vox spinning disk confocal system and a 60x Plan Apo VC oil immersion objective (NA 1.40). GFP (excitation: 488 nm, emission: 525 nm) and mCHERRY (excitation: 561 nm, emission: 615 nm) fluorescence were captured for 10-20 min at 30 frames.min^-1^.

The photokinesis unit of the same system was used for Fluorescence Recovery After Photobleaching (FRAP) experiments on *ralf22-2 x pRALF22::mCHERRY-RALF22_mature_* RHs. To this end, growing RHs were imaged for 3-4min after which mCHERRY was bleached in 3 ROIs corresponding to the growing tip, the shank just beneath the expanding dome and a region of the shank ∼100µm below the tip. Recovery of mCHERRY fluorescence was imaged for an additional 10min.

High resolution 3D imaging of mCHERRY-RALF22_mature_ localization in the RH CW was performed on a Nikon Ti2 W1-SoRa spinning disk system using the 561 nm laser line, 615 nm emission filter and a 100x SR HP Plan Apo silicon immersion objective (NA 1.35). Images were collected in xyz mode with a z-spacing of 0.25µm. 3D deconvolution was performed using NIS Elements AR, based on a calculated Point Spread Function (PSF) and 10-20 iterations.

3D acquisitions of *ralf22-2 x pRALF22::mCHERRY-RALF22_mature_* labelled with LM20, 2F4 or PAM1 antibodies were collected in the same way using the 561 nm and 488 nm laser lines respectively.

### Cloning

The *ralf22-1* mutant was generated using CRISPR-cas9 mutagenesis to induce a 182-nucleotide deletion downstream of the *RALF22* start codon. To this end, two domesticated guide RNA’s were simultaneously expressed (gRNA1: ATTGGCGATAGTAATCTCAGCCGGTTT; gRNA2: ATTgTAGCTACGGTGCTATGAGGGTTT).

Other constructs were generated by synthesizing the fragment of interest into pDONR221 (Thermofisher) and subsequent recombination to the desired destination vectors using gateway cloning. The *pRALF22::GFP* transgene was constructed using the entire 2658bp RALF22 promoter sequence, including the 5’-UTR. To generate the *pRALF22::RALF22* construct, the entire *RALF22* promoter and 360bp genomic sequence including STOP codon were synthesized. The *pRALF22::mCHERRY-RALF22_mature_*construct was generated by synthesizing the full *RALF22* genomic sequence in which mCHERRY was inserted between the sequences coding for the serine-protease cleavage site and the mature RALF22 peptide. To generate a RH-transcribed G5A [Ca^2+^] BRET sensor we synthesized a construct containing the 535bp promoter of the *EXPANSINa7* gene and the G5A sequence^54^. All pDONR221 plasmids were amplified in *E. coli*. Positive colonies were identified by PCR using *pRALF22*-(FW: TTCTCTGACGCCGTCGACTTTATC, REV: GGGTAGCACTATTTCTGCGTTGAC) and *GFP*-specific (FW: CACATGAAGCAGCACGACT, REV: TGCTCAGGTAGTGGTTGTCG) primers. Transgenes were recombined into pFAST-R01 (no tag, *pOLE1::RFP* seed marker), pFAST-R07 (C-terminal GFP, *pOLE1::RFP* seed marker) or pGWB1 (no tag, Kan/hyg resistance) using the gateway LR reaction^73^. *Agrobacterium tumefaciens* strain LBA4404 transformed with the construct of interest was used to transform Col-0 or *ralf22-2* plants using the floral dip method^74^. To generate the *pRALF22::RALF22^R82A,R90A,R^*^100^*^A^* construct, the full *pRALF22* promoter and *RALF22* coding sequence (CDS) were amplified from *Arabidopsis thaliana* Col-0 genomic DNA. Structure-based mutations were introduced in the *RALF22* CDS by site-directed mutagenesis (R82A,R90A,R100A). The transgene was recombined into pGGZ003 (addgene ID:48869) using GreenGate assembly and subsequently transformed into *Agrobacterium tumefaciens* strain ASE^75^.

### Image analysis

Image analysis was performed in Fiji. Root hair length was measured for 7 representative RHs per root, and >20 roots grown on 5 plates.

For *pRALF22::GFP* stacks, the 3D viewer and z-project plugins were used to generate 3D renderings and maximal projections of the acquired z-stacks.

To extract GCaMP3, FITC, TRITC and PI dynamics from timelapse acquisitions, individual frames of the red channel were aligned at subpixel resolution using the template matching plugin and normalized cross correlation as a matching method. The alignment coordinates were used to calculate the subpixel displacement of each frame relative to the first frame, from which the growth rate at each timepoint was calculated. The coordinates were then applied to the corresponding green channel. [Ca^2+^] traces were extracted from circular ROI in cytoplasm at the tip whereas the pH_ext_ was calculated from the FITC/TRITC ratio extracted from a ROI positioned in the extracellular medium along the expanding dome. The pH_ext_ was calibrated by imaging FITC and TRITC fluorescence in citric acid-sodium phosphate buffered liquid medium, at the exact same imaging settings. A ROI encompassing the apical CW was defined to extract the FITC and TRITC signal corresponding to a change in CW porosity or the PI signal corresponding to changes in CW charge. To quantify changes in cell width, the lateral displacement of the PI-stained CW just beneath the dome was calculated by laterally aligning the CW in a 1µm thin ROI (from the center of the cell to the frame’s edge). The alignment coordinates were used to calculate the radial expansion of the cell upon EGCG treatment.

To visualize the mobility of mCHERRY-RALF22_mature_ microdomains, we generated kymographs for a linear ROI overlaying the RH CW. To visualize microdomain mobility in the growing RH tip, a kymograph was generated from the apical CW which was straightened in Fiji. The recovery of mCHERRY-RALF22_mature_ fluorescence after bleaching was quantified by extracting the average fluorescence intensity over time in a ROI overlaying the stretch of bleached shank CW. Tip fluorescence recovery was quantified at the very apex after alignment of consecutive frames using the template matching plugin.

To investigate the dynamics of RALF22 secretion, mCHERRY-RALF22_mature_ fluorescence was quantified in a ROI at the very tip of frame aligned timelapse acquisitions, and correlated with the RH growth rate oscillations that were calculated from the alignment coordinates. RALF22 spacing in the CW was quantified by extracting the mCHERRY-RALF22_mature_ fluorescence intensity pattern from the line pattern present in mCHERRY-RALF22_mature_ kymographs or from a linear ROI drawn along the shank or tip RH CW. The obtained intensity data was detrended in AutoSignal 1.7 (Systat Software) by subtracting a cubic fit. Given that we were interested in the spatial frequency pattern represented by the spacing between individual microdomains, we used Fourier filtering to remove lower frequency oscillation (<0.6Hz) patterns associated with differences in fluorescence intensity along longer stretches of CW. We then generated a continuous wavelet time-frequency spectrum for each trace. The spatial frequency (mCHERRY-RALF22 fluorescence cycles.µm^-1^ -power distributions of all RHs were averaged and a gaussian distribution was fitted to get quantitative data on the mCHERRY-RALF22_mature_ microdomain/ring spacing. The ring spacing was calculated as such: ring spacing (µm)= 1 / spatial frequency (mCHERRY-fluorescence cycles.µm^-1^). Similarly, the predominant period characterizing the oscillatory growth rate pattern was extracted from the power distributions that described the continuous wavelet time-frequency spectrum of the growth rate oscillograms.

Based on the mCHERRY-RALF22_mature_, 2F4, LM20 and PAM1 Z-stacks, the lateral CW was reconstructed using the 3D project plugin. Colocalization was assessed using the Costes threshold regression method represented in the Coloc2 plugin with 50-100 randomisations^76^.

### Calcium assay

To quantify the systemic [Ca^2+^]_cyt_ response in trichoblast cell files upon RALF22 treatment, we generated Col-0 plants expressing the BRET-based GFP-aequorin Ca^2+^indicator G5A^54^ under the control of the RH-specific EXPANSINa7 promoter. Col-0 x pEXPa7::G5A plants were grown in liquid medium in 96-well microplates and incubated overnight in the presence of coelenterazine, as previously described^54^. Following injection of 200nM RALF22, RALF22^R82A,R90A,R100A^ or RALF22^Y75A,Y78A^ the luminescence was quantified for 20min using a TriStar LB 941 Multimode microplate reader equipped with automated well-by-well injection (Berthold Technologies). To investigate the importance of HG methylesterification, plants were preincubated with 32µM EGCG for 30min, 1h, 2h, 3h or overnight prior to RALF22 treatment. The [Ca^2+^] response was compared between treatments by extracting the area under the curve.

### Pectin and recombinant protein production

Synthetic RALF22_mature_ (AQKKYISYGAMRRNSVPCSRRGASYYNCQRGAQANPYSRGCSTITRCRR), RALF22^R82A,R90A,R100A^ (AQKKYISYGAMARNSVPCSARGASYYNCQAGAQANPYSRGCSTITRCRR) and RALF22^Y75A,Y78A^ (AQKKAISAGAMRRNSVPCSRRGASYYNCQRGAQANPYSRGCSTITRCRR) peptides were chemically synthesized (ProteoGenix, Schiltigheim, France).

LRX1-RALF22, FER_ecd_, ERU_ecd_ and LLG1 were produced in insect cells as previously described^29^. Codon-optimized synthetic genes for expression in *Spodoptera frugiperda* (Invitrogen, GeneArt), coding for *Arabidopsis thaliana* LLG1 (residues 24-144, At5g56170), ERU_ecd_ (residues 28-425, At5g61350), FER_ecd_ (residues 1-447, At3g51550), LRX2 (residues 1 to 385; At1g62440), LRX1 (residues 28-404, At1g12040) and RALF22_mature_ (residues 71-119, At3g05490) domains were cloned into a modified pFastBac vector (Geneva Biotech), with a native, 30K^77^ or BIP (native signal peptide of *Drosophila melanogaster*) signal peptide; and a TEV (tobacco etch virus protease) cleavable N- or C-terminal StrepII-9xHis tag. *Trichoplusia ni* Tnao38 cells^78,79^ were infected with LLG1, ERULUS, FERONIA or co-infected with LRX1-RALF22 or LRX2-RALF22 with a multiplicity of infection (MOI) of 3 and incubated 26 hours at 28°C and 48 hours at 22°C at 110 rpm. The secreted proteins were purified from the supernatant by sequential Ni^2+^ or StrepII affinity chromatography. Ni^2+^ (HisTrap excel, Cytiva) and StrepII (Strep-Tactin Superflow high capacity, IBA Lifesciences) columns were equilibrated in 25mM KPi pH 7.8, 500mM NaCl and 25mM Tris pH 8.0, 250mM NaCl, 1mM EDTA respectively. The tags were cleaved with His-tagged TEV protease at 4°C overnight and removed by a second Ni^2+^ affinity chromatography step. Proteins were further purified by SEC on a Superdex 200 Increase 10/300 GL column (Cytiva) equilibrated in 20mM citrate pH 5.0, 150mM NaCl. Proteins were concentrated using Amicon Ultra concentrators (Millipore, molecular weight cut-off 3,000, 10,000 and 30,000), and SDS-PAGE was used to assess the purity and integrity of the different proteins.

High and low methoxyl pectin samples were obtained from CPKelco (Denmark). The degree of pectin methylesterification was determined by Fourier transform infrared spectroscopy (FT-IR) using an IS50 spectrometer (ThermoFisher Scientific, Courtaboeuf, France). Pectin solutions were prepared in demineralised water at a concentration of 1 g.L^-1^, after which the pH was adjusted to pH 6.0, for complete ionization of the carboxylic acid groups. 100µL of solution were put on a CaF_2_ support and dried overnight at 35°C. The IR transmittance was measured at wavenumbers 1800 cm^−1^ to 800 cm^−1^ at a resolution of 2 cm^−1^. 200 scans were run per sample and averaged to obtain mean spectra. The spectra were baseline corrected, and preprocessed (OPUS and TheUnscrambler softwares). The ratio (X) of the absorption intensity of the bands around 1740cm^−1^ (carbonyl (C = O) stretching) and 1600–1630cm^−1^ (carboxylate (COO^−^) stretching) was fitted into a calibration equation (Y = 136.86X + 3.987 to calculate the degree of methylesterification (Y)^80^. The analysis was performed in duplicate.

The monosaccharide composition was determined by liquid-gas chromatography. Approx. 5mg of pectin powder was hydrolyzed in 2M H_2_SO_4_ for 2 hours at 100°C in the presence of inositol as internal standard. Sugars were then reduced, acetylated and analyzed as alditol acetates^81^ by liquid-gas chromatography (Perkin Elmer, Clarus 580, Shelton, USA) mounted with a DB-225 fused-silica capillary column (J&W Scientific, Folsorn, CA, USA, at 207°C with H_2_ as carrier gas). A standard solution containing the individual neutral monosaccharides (arabinose, rhamnose, glucose, xylose, galactose and mannose) was treated similarly for calculation of the monosaccharide recovery rates. The analysis was performed in triplicate. The results were expressed as anhydrous sugar to take into account their polysaccharide form in the starting sample. Uronic acids were quantified with the automated colorimetric *m*-hydroxybiphenyl method^82^. The analysis was performed in triplicate.

### Molecular interaction assays

We used microscale thermophoresis to identify potential receptor-RALF22 and RALF22-pectin interactions. FER_ecd_, ERU_ecd_ or LLG1, purified from insect cells, were labeled with the amine-reactive dye NT-647 Red-NHS (NanoTemper, Munich, Germany) according to the manufacturer’s instructions. Prior to interaction analysis, protein quality was assessed using a Tycho NT.6 (NanoTemper, Munich, Germany). Labeled protein was dissolved in MST buffer (50mM Tris-HCl pH 7.8, 150mM NaCl, 10mM MgCl_2_, 0.05% Tween-20). Wild-type or mutant RALF22 was titrated (starting at 100µM) against 20nM of LLG1, FERecd or ERUecd and loaded into premium coated Monolith NT.115 MST capillaries (NanoTemper, Munich, Germany). For the ternary LLG1-RALF22-FER_ecd_ complex, LLG1 (40nM) and wild-type or mutant RALF22 (5µM) were preincubated on ice for 15min and subsequently titrated against 5µM of FER_ecd_. For RALF22-pectin interactions, wild-type or mutant RALF22 (200µM) was titrated against 40nM Alexa Fluor 647-labeled OG7-13^47^. Following 10min incubation at RT, the capillaries were subjected to thermophoresis using a Monolith NT.115 (NanoTemper) at medium MST power and a LED excitation power of 80%. Data were withheld at a S/N >5 and the lack of protein aggregation or ligand-induced initial fluorescence changes. MST data were analyzed using MO. Affinity analysis v2.2.4 NT (NanoTemper).

### Quartz Crystal Microbalance with Dissipation Monitoring (QCM-D)

QCM-D was carried out with a “Q-Sense Analyser” (Biolin Scientific, Gothenburg, Sweden) using gold-coated SiO_2_ base sensors (QSX 301), spin-coated with 1% polyallylamine hydrochloride (PAH, Sigma-Aldrich, product number 283223) dissolved in water. Using a peristaltic pump a 0.1% pectin solution, in acetate buffer (10mM Na-Acetate pH 5.6, 10mM NaCl), was flown over the PAH layer over the period indicated by the first boxed area in Figs. 3D-F and 5H,I. After a washing step with buffer, protein at a concentration of 1µg/ml (wild-type or mutant RALF22) or 10µg/ml (LRX2-RALF22), was flown over the pectin layer for the period indicated by the second boxed area in Figs. 3D-F and 5H,I, followed by a washing step. Data presented are frequency (ΔF) and dissipation rate (ΔD) of the 3^rd^ harmonic resonant frequency of the base gold-coated sensors. Before reutilization, sensors were cleaned with a “Piranha” solution (7 vol 95% H_2_SO_4_ added to 3 vol 30% H_2_O_2_; respect the order!) using the following protocol: the quartz was incubated (4 in a holder) for 3-5min in Piranha solution. Next, the holder with the quartz was moved to two successive baths with MilliQ water. Then, each quartz was individually rinsed with MilliQ water and dried under a nitrogen stream. If traces still appeared on the surface during drying, rinsing and drying steps were repeated.

### Protein structure modelling

The crystal structure of the RALF4 mature peptide (residues 59 to 110; PDB: 6TME)^29^ was used to build a homology model of the mature RALF22 peptide (residues 71 to 119) through the Swiss-Model homology-modeling server (https://swissmodel.expasy.org/). The docking of LRX2 (PDB:6QXP) and the AlphaFold2 model of LRX1 with the homology model of RALF22 was performed using the Hawdock server^83^. The N-terminal region of RALF22 (74-88) was docked into the LLG2 structures (PDB: 6A5E), using the Flexpepdock web-server^84,85^. The FER receptor was then superimposed according to the crystal structure PDB: 6A5E.

### Statistics

Statistics were performed in R^86^. All data are represented as mean±SEM, and n indicates the number of independent replicates. Significance (Cl=0.05) was assessed by two-way analysis of variance (ANOVA, parametric) using linear mixed-effects models followed by a TukeyHSD (for pairwise statistical comparison), or by a Kruskal-Wallis test (non-parametric). Normality or deviations thereof were assessed with the Shapiro Wilkinson test (non-normal distribution < W=0.95< normal distribution) and by plotting the corresponding linear model’s residual values.

### movie legends

movie 1. RALF22 loss-of-function root hairs display aberrant growth and frequent bursting. Representative 2h timelapse acquisitions of growing Col-0 and *ralf22-2* (-/-) root hairs.

movie 2. Exogenous RALF22 supplementation induces a FERONIA-dependent growth arrest/signaling response and a FERONIA-independent increase in cell wall porosity. Representative longitudinal optical section timelapse acquisitions (10min) of growing Col-0 and *fer-4* RHs expressing the [Ca^2+^] sensor GCaMP3, in the presence of the dextran coupled dyes FITC (110kDa dextran, pH-sensitive) and TRITC (20kDa dextran, pH-insensitive). RHs were imaged in a microfluidics chip for 4min prior to addition of 5µM RALF22. The RHs response was followed for another 6 min (scale bar=5µm).

movie3. RALF22 is secreted to the root hair cell wall where it forms immobile periodic microdomains. (A) Representative longitudinal optical section timelapse acquisition (10min) of a growing *ralf22-2 x pRALF22::mCHERRY-RALF22_mature_* RH showing mCHERRY-RALF22_mature_ localization to the entire RH CW. Corresponding kymograph showing the immobility of secreted mCHERRY-RALF22mature microdomains (shown as vertical red lines) throughout the acquisition (scale bar=5µm). (B) RALF22 microdomains form in the growing RH dome. Consecutive timelapse frames of the growing tip have been aligned to allow visual tracking of mCHERRY-RALF22_mature_ microdomains as they move from tip to shank as the tip grows forward. The concave shape of the growing dome was straightened to generate a kymograph. Red lines in the kymograph depict mCHERRY-RALF22_mature_ microdomains, which originate at the very apex and remain immobile as they move towards the shank (scale bar=5µm).

movie 4. RALF22 is secreted at the growing root hair tip. Representative longitudinal optical section timelapse acquisition (10 min) of a growing *ralf22-2 x pRALF22::mCHERRY-RALF22_mature_* RH. After 4min of steady state growth, mCHERRY-RALF22mature fluorescence was bleached in a ROI in the tip and shank. Fluorescence Recovery After Photobleaching was followed for an additional 6min. Rapid mCHERRY-RALF22_mature_ fluorescence recovery was observed in the tip, but not in subapical regions (scale bar=5µm).

movie 5. The *lrx1-1/2-1* root hair phenotype is indistinguishable from *ralf22-2*. Representative 2h timelapse acquisitions of growing Col-0, *ralf22-2* and *lrx1-1/2-1* RHs.

movie 6. Pectin demethylesterification is crucial for root hair growth. Representative longitudinal optical section timelapse acquisitions (10 min) of growing Col-0 RHs expressing the [Ca^2+^] sensor GCaMP3 in the of dextran coupled FITC (110kDa dextran, pH-sensitive) and propidium iodide (CW charge reporter). Steady state RH growth was imaged for 4min prior to treatment with the Pectin Methylesterase (PME) inhibitor EGCG (100µM) or growth medium (mock). The RH’s response was followed for an additional 6min.

movie 7. LRX1/2 is required for RALF22 secretion. Representative longitudinal optical section timelapse acquisitions (10min) of growing *ralf22-2 x pRALF22::mCHERRY-RALF22_mature_* and *lrx1-1/2-1 x pRALF22::mCHERRY-RALF22_mature_* RHs illustrating the accumulation of mCHERRY-RALF22_mature_ in intracellular compartments in *lrx1-1/2-1* RHs.

## Notes

### Competing Interest Statement

The authors have declared no competing interest.

## References

1. Kjellén, L., & Lindahl, U. (2018). Specificity of glycosaminoglycan–protein interactions. Current Opinion in Structural Biology, 50, 101–108. 10.1016/j.sbi.2017.12.011

2. Haas, K. T., Wightman, R., Peaucelle, A., & Höfte, H. (2021). The role of pectin phase separation in plant cell wall assembly and growth. The Cell Surface, 7, 100054. 10.1016/j.tcsw.2021.100054

3. Coen, E., & Cosgrove, D. J. (2023). The mechanics of plant morphogenesis. Science (New York, N.Y.), 379(452), eade8055. 10.1126/science.ade8055

4. Wilson, L. A., Deligey, F., Wang, T., & Cosgrove, D. J. (2021). Saccharide analysis of onion outer epidermal walls. Biotechnology for Biofuels, 14(1), 1–14. 10.1186/s13068-021-01923-z

5. Levesque-Tremblay, G., Pelloux, J., Braybrook, S. A., & Müller, K. (2015). Tuning of pectin methylesterification: consequences for cell wall biomechanics and development. Planta, 242(4), 791–811. 10.1007/s00425-015-2358-5

6. Vincent, R. R., & Williams, M. A. K. (2009). Microrheological investigations give insights into the microstructure and functionality of pectin gels. Carbohydrate Research, 344(14), 1863– 1871. 10.1016/j.carres.2008.11.021

7. Derbyshire, P., McCann, M. C., & Roberts, K. (2007). Restricted cell elongation in Arabidopsis hypocotyls is associated with a reduced average pectin esterification level. BMC Plant Biology, 7(31), 1–12. 10.1186/1471-2229-7-31

8. Boyer, J. S. (2016). Enzyme-less growth in chara and terrestrial plants. Frontiers in Plant Science, 7(June), 1–15. 10.3389/fpls.2016.00866

9. Rojas, E. R., Hotton, S., & Dumais, J. (2011). Chemically mediated mechanical expansion of the pollen tube cell wall. Biophysical Journal, 101(8), 1844–1853. 10.1016/j.bpj.2011.08.016

10. Dumais, J. (2021). Mechanics and hydraulics of pollen tube growth. New Phytologist, 232(4), 1549–1565. 10.1111/nph.17722

11. Haas, K. T., Wightman, R., Meyerowitz, E. M., & Peaucelle, A. (2020). Pectin homogalacturonan nanofilament expansion drives morphogenesis in plant epidermal cells. Science, 367(6481), 1003–1007. 10.1126/science.aaz5103

12. Ma, Y., Jonsson, K., Aryal, B., De Veylder, L., Hamant, O., & Bhalerao, R. P. (2022). Endoreplication mediates cell size control via mechanochemical signaling from cell wall. Science Advances, 8(49), 1–11. 10.1126/sciadv.abq2047

13. Bidhendi, A. J., & Geitmann, A. (2016). Relating the mechanics of the primary plant cell wall to morphogenesis. Journal of Experimental Botany, 67(2), 449–461. 10.1093/jxb/erv535

14. Geitmann, A., & Ortega, J. K. E. (2009). Mechanics and modeling of plant cell growth. Trends in Plant Science, 14(9), 467–478. 10.1016/j.tplants.2009.07.006

15. Palin, R., & Geitmann, A. (2012). The role of pectin in plant morphogenesis. BioSystems, 109(3), 397–402. 10.1016/j.biosystems.2012.04.006

16. Richter, R. P., Baranova, N. S., Day, A. J., & Kwok, J. C. (2018). Glycosaminoglycans in extracellular matrix organisation: are concepts from soft matter physics key to understanding the formation of perineuronal nets? Current Opinion in Structural Biology, 50, 65–74. 10.1016/j.sbi.2017.12.002

17. Valentin, R., Cerclier, C., Geneix, N., Aguié-Béghin, V., Gaillard, C., Ralet, M. C., & Cathala, B. (2010). Elaboration of extensin-pectin thin film model of primary plant cell wall. Langmuir, 26(12), 9891–9898. 10.1021/la100265d

18. Fry, S. C. (1982). Isodityrosine, a new cross-linking amino acid from plant cell-wall glycoprotein. The Biochemical Journal, 204(2), 449–455. 10.1042/bj2040449 Brady, J. D., Sadler, I. H., & Fry, S. C. (1996). Di-isodityrosine, a novel tetrameric derivative of tyrosine in plant cell wall proteins: A new potential cross-link. Biochemical Journal, 315(1), 323–327. http://doi.org/10.1042/bj3150323

19. Held, M. A., Tan, L., Kamyabi, A., Hare, M., Shpak, E., & Kieliszewski, M. J. (2004). Di-isodityrosine is the intermolecular cross-link of isodityrosine-rich extensin analogs cross-linked in vitro. Journal of Biological Chemistry, 279(53), 55474–55482. 10.1074/jbc.M408396200

20. Moussu, S., & Ingram, G. (2023). The EXTENSIN enigma. The Cell Surface, 9, 100094. 10.1016/j.tcsw.2023.100094

21. Pearce, G., Moura, D. S., Stratmann, J., & Ryan, C. A. (2001). RALF, a 5-kDa ubiquitous polypeptide in plants, arrests root growth and development. Proceedings of the National Academy of Sciences of the United States of America, 98(22), 12843–12847. 10.1073/pnas.201416998

22. Abarca, A., Franck, C. M., & Zipfel, C. (2021). Family-wide evaluation of rapid alkalinization factor peptides. Plant Physiology, 187(2), 996–1010. 10.1093/plphys/kiab308

23. Xiao, Y., Stegmann, M., Han, Z., DeFalco, T. A., Parys, K., Xu, L., Belkhadir, Y., Zipfel, C., & Chai, J. (2019). Mechanisms of RALF peptide perception by a heterotypic receptor complex. Nature, 572(7768), 270–274. 10.1038/s41586-019-1409-7

24. Ge, Z., Bergonci, T., Zhao, Y., Zou, Y., Du, S., Liu, M.-C., Luo, X., Ruan, H., Garcia-Valencia, L. E., Zhong, S., Hou, S., Huang, Q., Lai, L., Moura, D. S., Gu, H., Dong, J., Wu, H.-M., Dresselhaus, T., Xiao, J., … Qu, L.-J. (2017). Arabidopsis pollen tube integrity and sperm release are regulated by RALF-mediated signaling. Science, 358(December), 1596–1600. 10.1126/science.aao3642

25. Ge, Z., Zhao, Y., Liu, M. C., Zhou, L. Z., Wang, L., Zhong, S., Hou, S., Jiang, J., Liu, T., Huang, Q., Xiao, J., Gu, H., Wu, H. M., Dong, J., Dresselhaus, T., Cheung, A. Y., & Qu, L. J. (2019). LLG2/3 Are Co-receptors in BUPS/ANX-RALF Signaling to Regulate Arabidopsis Pollen Tube Integrity. Current Biology, 29(19), 3256–3265.e5. 10.1016/j.cub.2019.08.032

26. Gao, Q., Wang, C., Xi, Y., Shao, Q., Hou, C., Li, L., & Luan, S. (2023). RALF signaling pathway activates MLO calcium channels to maintain pollen tube integrity. Cell Research, 33(1), 71– 79. 10.1038/s41422-022-00754-3

27. Mecchia, M. A., Santos-Fernandez, G., Duss, N. N., Somoza, S. C., Boisson-Dernier, A., Gagliardini, V., Martínez-Bernardini, A., Fabrice, T. N., Ringli, C., Muschietti, J. P., & Grossniklaus, U. (2017). RALF4/19 peptides interact with LRX proteins to control pollen tube growth in Arabidopsis. Science, 358(6370), 1600–1603. 10.1126/science.aao5467

28. Solis-Miranda, J., & Quinto, C. (2021). The CrRLK1L subfamily: One of the keys to versatility in plants. Plant Physiology and Biochemistry, 166(May), 88–102. 10.1016/j.plaphy.2021.05.028

29. Moussu, S., Broyart, C., Santos-Fernandez, G., Augustin, S., Wehrle, S., Grossniklaus, U., & Santiago, J. (2020). Structural basis for recognition of RALF peptides by LRX proteins during pollen tube growth. Proceedings of the National Academy of Sciences of the United States of America, 117(13), 7494–7503. 10.1073/pnas.2000100117

30. Herger, A., Gupta, S., Kadler, G., Franck, C. M., Boisson-Dernier, A., & Ringli, C. (2020). Overlapping functions and protein-protein interactions of LRR-extensins in Arabidopsis. PLoS Genetics, 16(6), 1–24. 10.1371/journal.p gen.1008847

31. Zhao, C., Zayed, O., Yu, Z., Jiang, W., Zhu, P., Hsu, C. C., Zhang, L., Andy Tao, W., Lozano-Durán, R., & Zhu, J. K. (2018). Leucine-rich repeat extensin proteins regulate plant salt tolerance in Arabidopsis. Proceedings of the National Academy of Sciences of the United States of America, 115(51), 13123–13128. 10.1073/pnas.1816991115

32. Baumberger, N., Doesseger, B., Guyot, R., Diet, A., Parsons, R. L., Clark, M. A., Simmons, M. P., Bedinger, P., Goff, S. A., Ringli, C., & Keller, B. (2003). Whole-genome comparison of leucine-rich repeat extensins in Arabidopsis and rice. A conserved family of cell wall proteins form a vegetative and a reproductive clade. Plant Physiology, 131(3), 1313–1326. 10.1104/pp.102.014928

33. Schoenaers, S., Balcerowicz, D., & Vissenberg, K. (2017). Molecular Mechanisms Regulating Root Hair Tip Growth: A Comparison with Pollen Tubes. In Pollen Tip Growth: From Biophysical Aspects to Systems Biology (pp. 167–243). Springer. 10.1007/978-3-319-56645-0_9

34. Hocq, L., Pelloux, J., & Lefebvre, V. (2017). Connecting Homogalacturonan-Type Pectin Remodeling to Acid Growth. Trends in Plant Science, 22(1), 20–29. 10.1016/j.tplants.2016.10.009

35. Pacifici, E., Di Mambro, R., Dello Ioio, R., Costantino, P., & Sabatini, S. (2018). Acidic cell elongation drives cell differentiation in the Arabidopsis root. The EMBO Journal, 37(16), 1–9. 10.15252/embj.201899134

36. Denyer, T., Ma, X., Klesen, S., Scacchi, E., Nieselt, K., & Timmermans, M. C. P. (2019). Spatiotemporal Developmental Trajectories in the Arabidopsis Root Revealed Using High-Throughput Single-Cell RNA Sequencing. Developmental Cell, 48(6), 840–852.e5. 10.1016/j.devcel.2019.02.022

37. Stegmann, M., Monaghan, J., Smakowska-Luzan, E., Rovenich, H., Lehner, A., Holton, N., Belkhadir, Y., & Zipfel, C. (2017). The receptor kinase FER is a RALF-regulated scaffold controlling plant immune signaling. Science, 355(6322), 287–289. 10.1126/science.aal2541

38. Gonneau, M., Desprez, T., Martin, M., Doblas, V. G., Bacete, L., Miart, F., Sormani, R., Hématy, K., Renou, J., Landrein, B., Murphy, E., Van De Cotte, B., Vernhettes, S., De Smet, I., & Höfte, H. (2018). Receptor Kinase THESEUS1 Is a Rapid Alkalinization Factor 34 Receptor in Arabidopsis. Current Biology, 2452–2458. 10.1016/j.cub.2018.05.075

39. Srivastava, R., Liu, J. X., Guo, H., Yin, Y., & Howell, S. H. (2009). Regulation and processing of a plant peptide hormone, AtRALF23, in Arabidopsis. Plant Journal, 59(6), 930–939. 10.1111/j.1365-313X.2009.03926.x

40. Zhao, C., Zayed, O., Yu, Z., Jiang, W., Zhu, P., Hsu, C. C., Zhang, L., Andy Tao, W., Lozano-Durán, R., & Zhu, J. K. (2018). Leucine-rich repeat extensin proteins regulate plant salt tolerance in Arabidopsis. Proceedings of the National Academy of Sciences of the United States of America, 115(51), 13123–13128. 10.1073/pnas.1816991115

41. Schoenaers, S., Balcerowicz, D., Breen, G., Hill, K., Zdanio, M., Mouille, G., Holman, T. J., Oh, J., Wilson, M. H., Nikonorova, N., Vu, L. D., De Smet, I., Swarup, R., De Vos, W. H., Pintelon, I., Adriaensen, D., Grierson, C., Bennett, M. J., & Vissenberg, K. (2018). The Auxin-Regulated CrRLK1L Kinase ERULUS Controls Cell Wall Composition during Root Hair Tip Growth. Current Biology, 28(5), 722–732.e6. 10.1016/j.cub.2018.01.050

42. Duan, Q., Kita, D., Li, C., Cheung, A. Y., & Wu, H. M. (2010). FERONIA receptor-like kinase regulates RHO GTPase signaling of root hair development. Proceedings of the National Academy of Sciences of the United States of America, 107(41), 17821–17826. 10.1073/pnas.1005366107

43. Shen, Q., Bourdais, G., Pan, H., Robatzek, S., & Tang, Di. (2017). Arabidopsis glycosylphosphatidylinositol-anchored protein LLG1 associates with and modulates FLS2 to regulate innate immunity. Proceedings of the National Academy of Sciences of the United States of America, 114(22), 5749–5754. 10.1073/pnas.1614468114

44. Toyota, M., Spencer, D., Sawai-toyota, S., Jiaqi, W., Zhang, T., Koo, A. J., Howe, G. A., & Gilroy, S. (2018). Glutamate triggers long-distance, calcium-based plant defense signaling. Science, 361(September), 1112–1115. 10.1126/science.aat7744

45. Voetmann, L. M., Rolin, B., Kirk, R. K., Pyke, C., & Hansen, A. K. (2023). The intestinal permeability marker FITC-dextran 4kDa should be dosed according to lean body mass in obese mice. Nutrition and Diabetes, 13(1), 4–8. 10.1038/s41387-022-00230-2

46. Chen, H., Liu, Y.-C., Zhang, Z., Li, M., Du, L., Wu, P.-C., Chong, W.-H., Ren, F., Zheng, W., & Liu, T.-M. (2022). Mouse Strain– and Charge-Dependent Vessel Permeability of Nanoparticles at the Lower Size Limit. Frontiers in Chemistry, 10(July), 1–9. 10.3389/fchem.2022.944556

47. Mravec, J., Kračun, S. K., Rydahl, M. G., Westereng, B., Pontiggia, D., De Lorenzo, G., Domozych, D. S., & Willats, W. G. T. (2017). An oligogalacturonide-derived molecular probe demonstrates the dynamics of calcium-mediated pectin complexation in cell walls of tip-growing structures. Plant Journal, 91(3), 534–546. 10.1111/tpj.13574

48. Jaafar, Z., Mazeau, K., Boissière, A., Le Gall, S., Villares, A., Vigouroux, J., Beury, N., Moreau, C., Lahaye, M., & Cathala, B. (2019). Meaning of xylan acetylation on xylan-cellulose interactions: A quartz crystal microbalance with dissipation (QCM-D) and molecular dynamic study. Carbohydrate Polymers, 226(September), 115315. 10.1016/j.carbpol.2019.115315

49. Tanaka, T. (1981). Gels. Scientific American, 124–138. 10.1038/scientificamerican0181-124

50. Shibayama, M., & Tanaka, T. (1993). Volume phase transition and related phenomena of polymer gels. Advances in Polymer Science, 109, 1–60. 10.1007/3-540-56791-7_1

51. Baumberger, N., Steiner, M., Ryser, U., Keller, B., & Ringli, C. (2003). Synergistic interaction of the two paralogous Arabidopsis genes LRX1 and LRX2 in cell wall formation during root hair development. Plant Journal, 35(1), 71–81. 10.1046/j.1365-313X.2003.01784.x

52. Baumberger, N., Ringli, C., & Keller, B. (2001). The chimeric leucine-rich repeat/extensin cell wall protein LRX1 is required for root hair morphogenesis in Arabidopsis thaliana. Genes and Development, 15(9), 1128–1139. 10.1101/gad.200201

53. Rounds, C. M., Lubeck, E., Hepler, P. K., & Winship, L. J. (2011). Propidium iodide competes with Ca^2+^ to label pectin in pollen tubes and Arabidopsis root hairs. Plant Physiology, 157(1), 175–187. 10.1104/pp.111.182196

54. Xiong, T. C., Ronzier, E., Sanchez, F., Corratgé-Faillie, C., Mazars, C., & Thibaud, J. B. (2014). Imaging long distance propagating calcium signals in intact plant leaves with the BRET-based GFP-aequorin reporter. Frontiers in Plant Science, 5(43), 1–13. 10.3389/fpls.2014.00043

55. Xu, F., Gonneau, M., Faucher, E., Habrylo, O., Lefebvre, V., Domon, J. M., Martin, M., Sénéchal, F., Peaucelle, A., Pelloux, J., & Höfte, H. (2022). Biochemical characterization of Pectin Methylesterase Inhibitor 3 from Arabidopsis thaliana. The Cell Surface, 8(July), 4–11. 10.1016/j.tcsw.2022.100080

56. Verhertbruggen, Y., Marcus, S. E., Haeger, A., Ordaz-Ortiz, J. J., & Knox, J. P. (2009). An extended set of monoclonal antibodies to pectic homogalacturonan. Carbohydrate Research, 344(14), 1858–1862. 10.1016/j.carres.2008.11.010

57. Liners, F., Letesson, J. J., Didembourg, C., & Van Cutsem, P. (1989). Monoclonal Antibodies against Pectin: Recognition of a Conformation Induced by Calcium. Plant Physiology, 91(4), 1419–1424. 10.1104/pp.91.4.1419

58. Willats, W. G. T., Gilmartin, P. M., Mikkelsen, J. D., & Knox, J. P. (1999). Cell wall antibodies without immunization: Generation and use of de-esterified homogalacturonan block-specific antibodies from a naive phage display library. Plant Journal, 18(1), 57–65. 10.1046/j.1365-313X.1999.00427.x

59. Langford-Smith, A., Keenan, T. D. L., Clark, S. J., Bishop, P. N., & Day, A. J. (2014). The role of complement in age-related macular degeneration: Heparan sulphate, a ZIP code for complement factor H? Journal of Innate Immunity, 6(4), 407–416. 10.1159/000356513

60. Voxeur, A., Habrylo, O., Guénin, S., Miart, F., Soulié, M. C., Rihouey, C., Pau-Roblot, C., Domon, J. M., Gutierrez, L., Pelloux, J., Mouille, G., Fagard, M., Höfte, H., & Vernhettes, S. (2019). Oligogalacturonide production upon Arabidopsis thaliana-Botrytis cinerea interaction. Proceedings of the National Academy of Sciences of the United States of America, 116(39), 19743–19752. 10.1073/pnas.1900317116

61. Willats, W. G. T., McCartney, L., Steele-King, C. G., Marcus, S. E., Mort, A., Huisman, M., Van Alebeek, G. J., Schols, H. A., Voragen, A. G. J., Le Goff, A., Bonnin, E., Thibault, J. F., & Knox, J. P. (2004). A xylogalacturonan epitope is specifically associated with plant cell detachment. Planta, 218(4), 673–681. 10.1007/s00425-003-1147-8

62. Tanaka, T., Annaka, M., Ilmain, F., Ishii, K., Kokufuta, E., Suzuki, A., & Tokita, M. (1992). Phase Transitions of Gels. In NATO ASI Series, Vol. 64. Mechanics of Swelling (pp. 683–703). 10.1146/ANNUREV.MS.22.080192.001331

63. Cannon, M. C., Terneus, K., Hall, Q., Tan, L., Wang, Y., Wegenhart, B. L., Chen, L., Lamport, D. T. A., Chen, Y., & Kieliszewski, M. J. (2008). Self-assembly of the plant cell wall requires an extensin scaffold. Proceedings of the National Academy of Sciences of the United States of America, 105(6), 2226–2231. 10.1073/pnas.0711980105

64. Shih, H. W., Miller, N. D., Dai, C., Spalding, E. P., & Monshausen, G. B. (2014). The receptor-like kinase FERONIA is required for mechanical signal transduction in Arabidopsis seedlings. Current Biology, 24(16), 1887–1892. 10.1016/j.cub.2014.06.064

65. Tang, W., Lin, W., Zhou, X., Guo, J., Dang, X., Li, B., Lin, D., & Yang, Z. (2022). Mechano-transduction via the pectin-FERONIA complex activates ROP6 GTPase signaling in Arabidopsis pavement cell morphogenesis. Current Biology, 32(3), 508–517.e3. 10.1016/j.cub.2021.11.031

66. Feng, W., Kita, D., Peaucelle, A., Cartwright, H. N., Doan, V., Duan, Q., Liu, M. C., Maman, J., Steinhorst, L., Schmitz-Thom, I., Yvon, R., Kudla, J., Wu, H. M., Cheung, A. Y., & Dinneny, J. R. (2018). The FERONIA Receptor Kinase Maintains Cell-Wall Integrity during Salt Stress through Ca^2+^ Signaling. Current Biology, 28(5), 666–675.e5. 10.1016/j.cub.2018.01.023

67. Brost, C., Studtrucker, T., Reimann, R., Denninger, P., Czekalla, J., Krebs, M., Fabry, B., Schumacher, K., Grossmann, G., & Dietrich, P. (2019). Multiple cyclic nucleotide-gated channels coordinate calcium oscillations and polar growth of root hairs. Plant Journal, 99(5), 910–923. 10.1111/tpj.14371

68. Park, S., Szumlanski, A. L., Gu, F., Guo, F., & Nielsen, E. (2011). A role for CSLD3 during cell-wall synthesis in apical plasma membranes of tip-growing root-hair cells. Nature Cell Biology, 13, 973–980.

49. Cavalier, D. M., Lerouxel, O., Neumetzler, L., Yamauchi, K., Reinecke, A., Freshour, G., Zabotina, O. A., Hahn, M. G., Burgert, I., Pauly, M., Raikhel, N. V., & Keegstra, K. (2008). Disrupting two Arabidopsis thaliana xylosyltransferase genes results in plants deficient in xyloglucan, a major primary cell wall component. Plant Cell, 20(6), 1519–1537. 10.1105/tpc.108.059873

70. Lin, C., Choi, H.-S., & Cho, H.-T. (2011). Root hair-specific EXPANSIN A7 is required for root hair elongation in Arabidopsis. Molecules and Cells, 31, 393–397. 10.1007/s10059-011-0046-2

71. Winter, D., Vinegar, B., Nahal, H., Ammar, R., Wilson, G. V., & Provart, N. J. (2007). An “electronic fluorescent pictograph” Browser for exploring and analyzing large-scale biological data sets. PLoS ONE, 2(8), 1–12. 10.1371/journal.pone.0000718

72. Herburger, K., Schoenaers, S., Vissenberg, K., & Mravec, J. (2022). Shank-localized cell wall growth contributes to Arabidopsis root hair elongation. Nature Plants, 8(11), 1222–1232. 10.1038/s41477-022-01259-y

73. Shimada, T. L., Shimada, T., & Hara-Nishimura, I. (2010). A rapid and non-destructive screenable marker, FAST, for identifying transformed seeds of Arabidopsis thaliana: TECHNICAL ADVANCE. Plant Journal, 61(3), 519–528. 10.1111/j.1365-313X.2009.04060.x

74. Clough, S. J., & Bent, A. F. (1998). Floral dip: A simplified method for Agrobacterium-mediated transformation of Arabidopsis thaliana. Plant Journal, 16(6), 735–743. 10.1046/j.1365-313X.1998.00343.x

75. Lampropoulos, A., Sutikovic, Z., Wenzl, C., Maegele, I., Lohmann, J. U., & Forner, J. (2013). GreenGate - A novel, versatile, and efficient cloning system for plant transgenesis. PLoS ONE, 8(12). 10.1371/journal.pone.0083043

76. Costes, S. V., Daelemans, D., Cho, E. H., Dobbin, Z., Pavlakis, G., & Lockett, S. (2004). Automatic and quantitative measurement of protein-protein colocalization in live cells. Biophysical Journal, 86(6), 3993–4003. 10.1529/biophysj.103.038422

77. Futatsumori-Sugai, M., & Tsumoto, K. (2010). Signal peptide design for improving recombinant protein secretion in the Baculovirus expression vector system. Biochemical and Biophysical Research Communications, 391(1), 931–935. 10.1016/j.bbrc.2009.11.167

78. Hashimoto, Y., Zhang, S., & Blissard, G. W. (2010). Ao38, a new cell line from eggs of the black witch moth, Ascalapha odorata (Lepidopteralll: Noctuidae), is permissive for AcMNPV infection and produces high levels of recombinant proteins. BMC Biotechnology, 10(50). 10.1186/1472-6750-10-50

79. Hashimoto, Y., Zhang, S., Zhang, S., Chen, Y. R., & Blissard, G. W. (2012). Correction: BTI-Tnao38, a new cell line derived from Trichoplusia ni, is permissive for AcMNPV infection and produces high levels of recombinant proteins. BMC Biotechnology, 12, 3–6. 10.1186/1472-6750-12-12

80. Kyomugasho, C., Christiaens, S., Shpigelman, A., Van Loey, A. M., & Hendrickx, M. E. (2015). FT-IR spectroscopy, a reliable method for routine analysis of the degree of methylesterification of pectin in different fruit- and vegetable-based matrices. Food Chemistry, 176, 82–90. 10.1016/j.foodchem.2014.12.033

81. Englyst, H., Wiggins, H. S., & Cummings, J. H. (1982). Determination of the non-starch polysaccharides in plant foods by gas - Liquid chromatography of constituent sugars as alditol acetates. The Analyst, 107(1272), 307–318. 10.1039/AN9820700307

82. Thibault, J.-F. (1979). Automatisation du dosage des substances pectiques par la méthode au méta-hidroxydiphenyl. Lebensmittel-Wissenschaft Und Technologie, 12(5), 247–251.

83. Weng, G., Wang, E., Wang, Z., Liu, H., Zhu, F., Li, D., & Hou, T. (2019). HawkDock: a web server to predict and analyze the protein-protein complex based on computational docking and MM/GBSA. Nucleic Acids Research, 47(W1), W322–W330. 10.1093/nar/gkz397

84. Raveh, B., London, N., & Schueler-Furman, O. (2010). Sub-angstrom modeling of complexes between flexible peptides and globular proteins. Proteins: Structure, Function and Bioinformatics, 78(9), 2029–2040. 10.1002/prot.22716

85. London, N., Raveh, B., Cohen, E., Fathi, G., & Schueler-Furman, O. (2011). Rosetta FlexPepDock web server - High resolution modeling of peptide-protein interactions. Nucleic Acids Research, 39(SUPPL. 2), 249–253. 10.1093/nar/gkr431

86. 86. Team, R. C. (2021). R: A language and environment for statistical computing. R Foundation for Statistical Computing, Vienna, Austria. https://www.r-project.org/

