## Supplemental Figures for "A pectin-binding peptide with a structural and signaling role in the assembly of the plant cell wall"

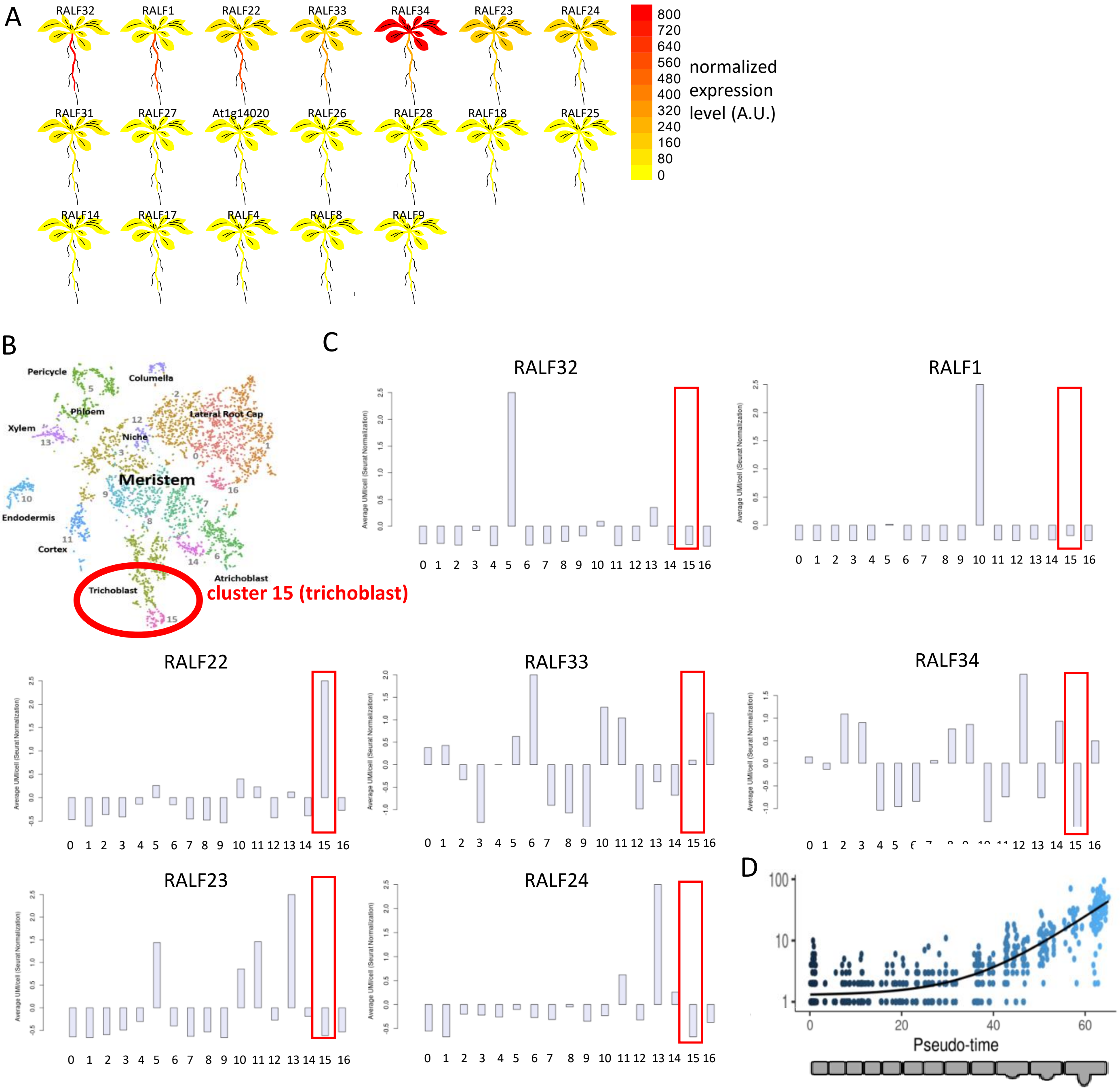


Fig. S1: Identification of trichoblast-expressed RALFs from public transcriptome data. (A) The relative expression for each RALF that is represented on the Affymetrix ATH1 array is shown, sorted by degree of expression in the primary root. (B) Annotated t-SNE plot showing the different cell types represented by the single cell transcriptome dataset (Denyer et al., 2019). Trichoblast cells are represented in cluster 15. (C) Seurat-normalized expression values for each cluster. The expression values corresponding to cluster 15 are highlighted in red. *RALF22* expression is enriched in trichoblast cells. (D) trichoblast pseudotime expression profile displaying *RALF22* expression in function of the trichoblast's distance to the meristem (i.e. its developmental stage).

A

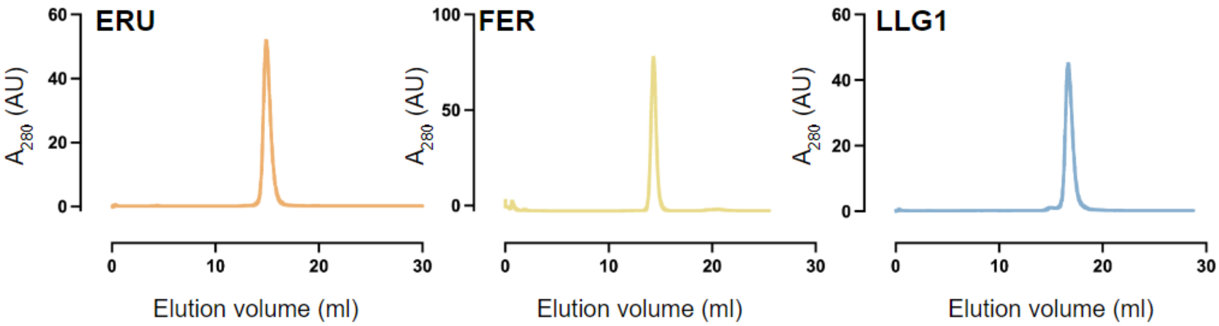


B

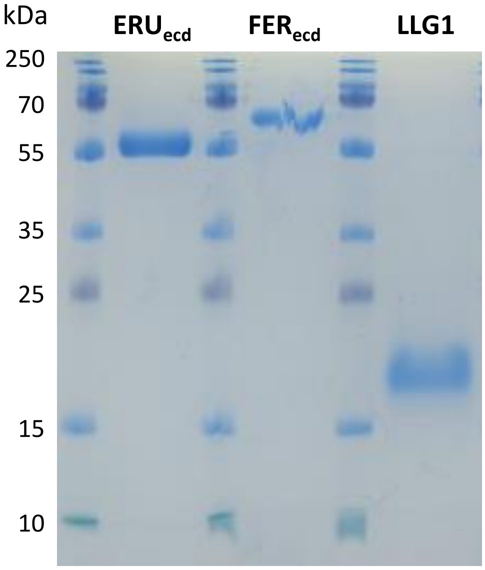


| | protein size (kDa) | N-linked glycosylation sites |
|---|---|---|
| ERUecd | 43.29 | 6 |
| FERecd | 48.81 | 10 |
| LLG1 | 13.32 | 1 |

C

| labelled protein | titrated protein | Kd |
|---|---|---|
| LLG1* | RALF22 | 6.09 ± 1.13 µM |
| LLG1* | RALF22$^{Y75A,Y78A}$ | 28.48 ± 2.43 µM |
| LLG1* | RALF22$^{R82A,R90A,R100A}$ | 2.95 ± 0.65 µM |
| LLG1* | FER$_{ecd}$ | no interaction |
| LLG1* | ERU$_{ecd}$ | no interaction |
| FER$_{ecd}$* | RALF22 | no interaction |
| ERU$_{ecd}$* | RALF22 | no interaction |
| LLG1*+RALF22 | FER$_{ecd}$ | 118.02 ± 59.01 nM |
| LLG1*+RALF22$^{Y75A,Y78A}$ | FER$_{ecd}$ | no interaction |
| LLG1*+RALF22 | ERU$_{ecd}$ | no interaction |
| LLG1*+RALF22$^{Y75A,Y78A}$ | ERU$_{ecd}$ | no interaction |
| FER$_{ecd}$* | ERU$_{ecd}$ | no interaction |
| ERU$_{ecd}$* | FER$_{ecd}$ | no interaction |
| ERU$_{ecd}$* | FER$_{ecd}$+(LLG1+RALF22) | no interaction |
| OG7-13* | RALF22 | 3.03 ± 0.53 µM |
| OG7-13* | RALF22$^{R82A,R90A,R100A}$ | 11.1 ± 2.65 µM |

Fig. S2. Characterization of recombinant protein extracts used for *in vitro* interaction analysis. (A) Analytical size-exclusion chromatography (SEC) traces of ERU$_{ecd}$, FER$_{ecd}$ and LLG1 purified from insect cells. (B) SDS-PAGE of the corresponding SEC protein fractions. Protein size is affected by N-linked glycosylation for ERU$_{ecd}$, FER$_{ecd}$ and LLG1. (C) Overview of the protein-protein and protein-oligosaccharide combinations for which binding affinities were determined using MicroScale Thermophoresis (MST). A potential interaction partner was titrated against a lysine-labeled protein or protein-complex target (*). The average Kd (n≥3) is reported for valid interactions.

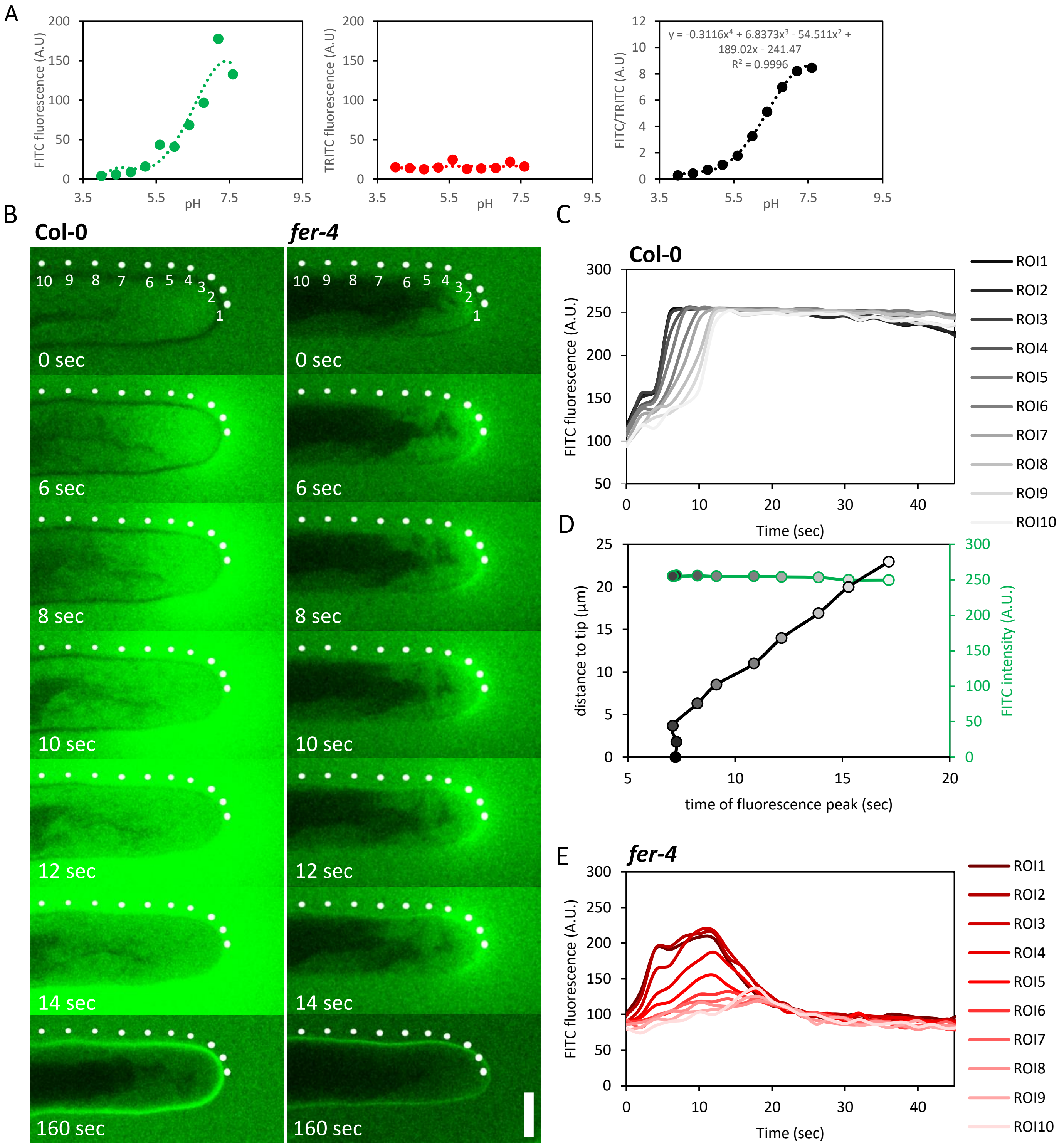


Fig. S3. Visualization and quantification of extracellular pH ($pH_{ex}$) dynamics using FITC-110kDa dextran and TRITC-20kDa dextran fluorescence. (A) calibration of FITC-110kDa dextran and TRITC-20kDa dextran fluorescence for ratiometric determination of the $pH_{ex}$. The 4th degree polynomal fit for the pH-dependent FITC/TRITC ratio was used for calculation of the $pH_{ext}$ dynamics during root hair growth. The intensity of both fluorophores was quantified in citric acid/sodium phosphate buffered minimal medium (pH 4.0-7.6) in microfluidics chips with the same imaging parameters used for confocal live cell imaging. Datapoints represent mean fluorescence ± SEM (n=3). (B-E) Tip-to-shank propagation of the extracellular alkalinization induced by RALF22 treatment depends on FERONIA. (B) consecutive frames of a representative growing root hair treated with 5µM RALF22, showing how the alkalinization that starts at the tip propagates towards the shank in Col-0 but not in *fer-4*. Dots indicate the ROIs used for fluorescence intensity quantification at different positions relative to the tip. (C) quantification of the extracellular FITC fluorescence in time, in Col-0, in the ROIs depicted in A. (D) Determination of the timing associated with maximum FITC fluorescence in each ROI in Col-0. (E) Quantification of the extracellular FITC fluorescence in each ROI in *fer-4*.

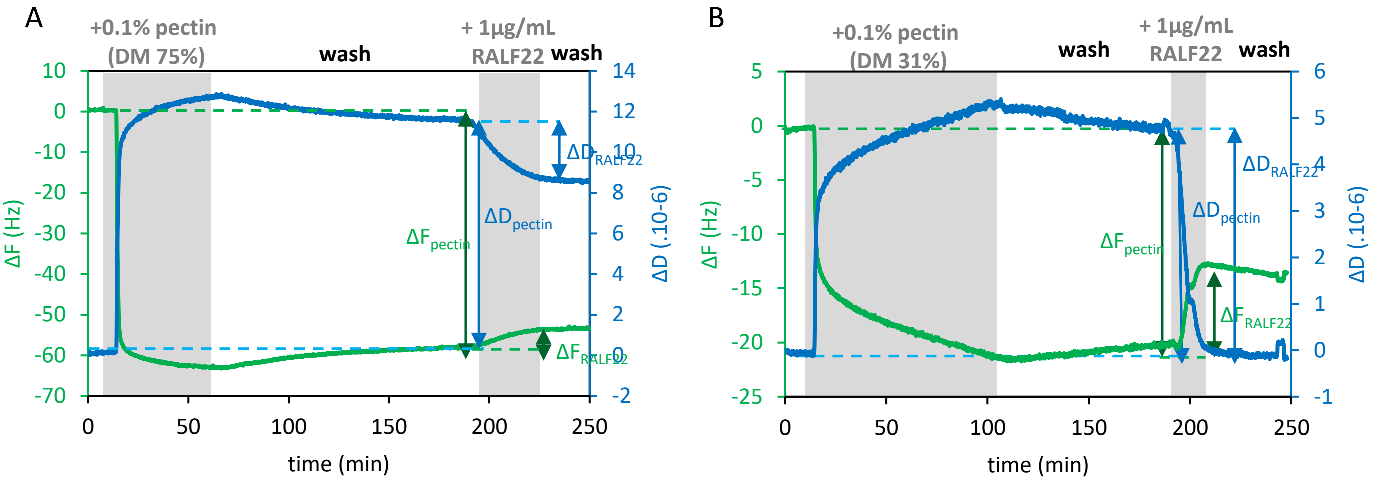


ΔF ($\Delta F_{RALF22}/\Delta F_{pectin}.100$)=8%
ΔD ($\Delta D_{RALF22}/\Delta D_{pectin}.100$)=25%

ΔF ($\Delta F_{RALF22}/\Delta F_{pectin}.100$)=30%
ΔD ($\Delta D_{RALF22}/\Delta D_{pectin}.100$)=94%

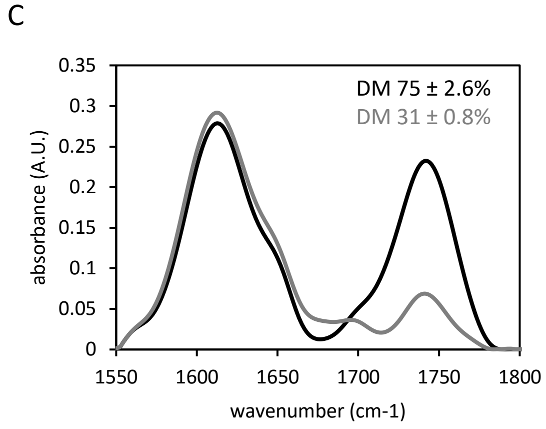


D

| Sugar (%) | Pectin DM 75% | | Pectin DM 31% | |
|---|---|---|---|---|
| Rhamnose | 3.00 | +/- 0,15 | 0.33 | +/- 0,05 |
| Fucose | - | - | - | - |
| Arabinose | 2.96 | +/- 0,15 | 0.32 | +/- 0,05 |
| Xylose | 0.39 | +/- 0,22 | 0.28 | +/- 0,15 |
| Mannose | - | - | - | - |
| Galactose | 4.97 | +/- 0,29 | 2.71 | +/- 0,12 |
| Glucose | 0.23 | +/- 0,06 | 0.33 | +/- 0,10 |
| Gal. A. | 80.62 | +/- 12,5 | 86.31 | +/- 5,5 |
| Total | 92.18 | +/- 12,99 | 90.29 | +/- 5,77 |

Fig. S4. QCM-D to monitor RALF22-induced changes in the mass and viscoelasticity of pectin layers with different degrees of methylesterification (DM). (A,B) Typical QCM-D experiments showing third harmonics of ΔF (green) and ΔD (blue) for the binding of a 0.1% pectin solution with a DM of 75% (A) and 31% (B) to the PAH layer, followed by a RALF22 (1µg/ml) solution. Grey zones depict the period during which pectin and RALF22 solutions are delivered. (C-D) characterization of the pectin solutions used for QCM-D analysis. (C) representative FT-IR spectra of DM 75% (black) and DM 31% (grey) pectin preparations showing the absorbance at wavenumber 1740 $cm^{-1}$ (ester carbonyl group stretching) and 1630-1600 $cm^{-1}$ (carboxylate group). The DM was calculated from the ratio between the absorbance at 1740 $cm^{-1}$ and the combined absorbance at 1740 and 1630-1600 $cm^{-1}$. (D) monosaccharide composition of DM 75% and DM 31% pectin preparations (n=3). Values represent mean ± SEM (n=3).

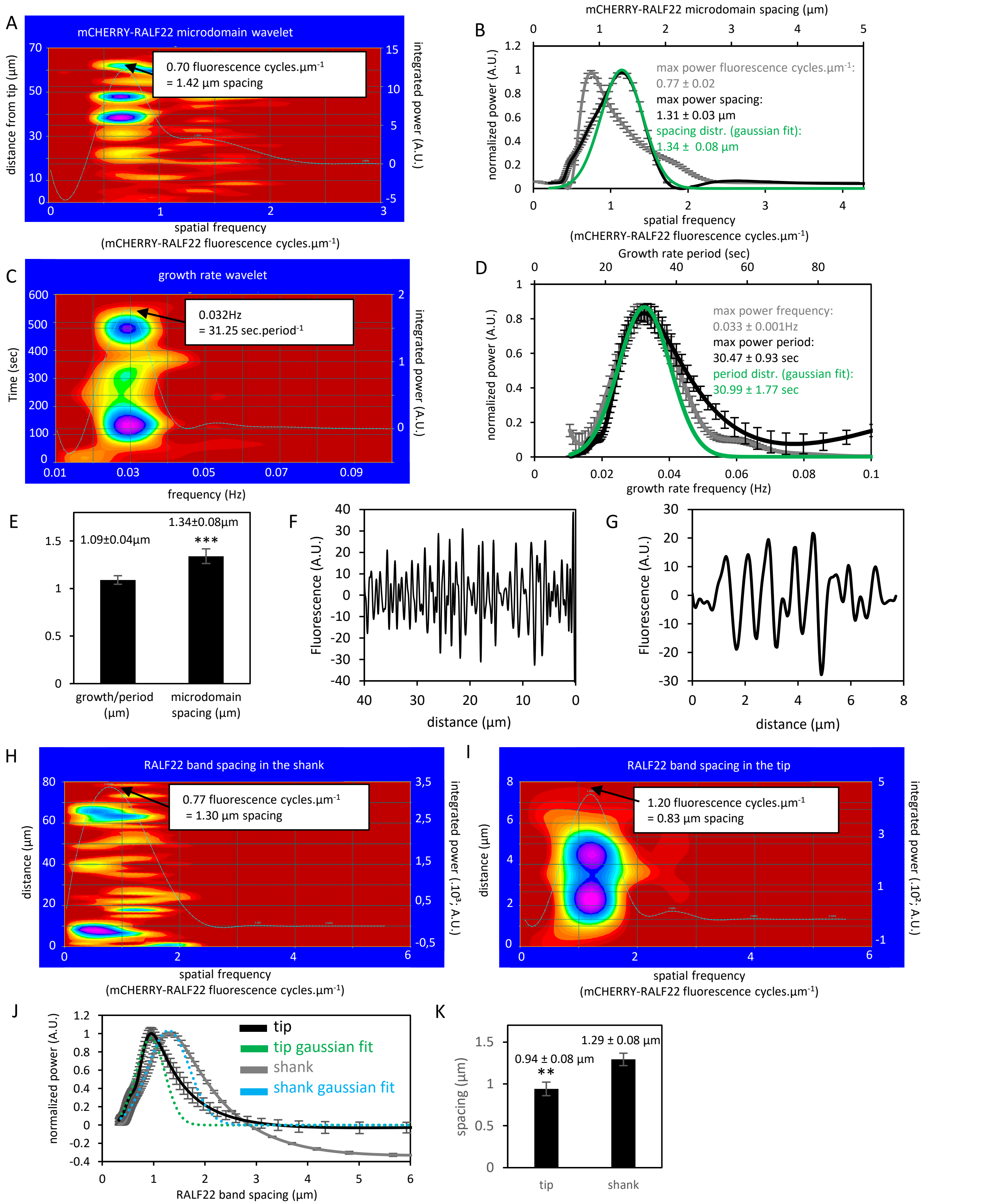


Fig. S5. Comparative analysis of the periodicity of RALF22 microdomains. (A) representative continuous wavelet spectrum showing the cumulative prevalence of spatial frequencies (power spectrum; white line) describing mCHERRY-RALF22$_{mature}$ fluorescence along the RH CW. (B) Corresponding average power spectra depicting the spatial frequency (grey) and microdomain spacing (µm; black) distributions along the RHs longitudinal axis. (C) representative continuous wavelet spectrum showing the cumulative prevalence of frequencies (0-0.1Hz, power spectrum; white line) describing an oscillatory growth rate trace. (D) average power spectra depicting the frequency (Hz; grey) and period (sec; black) distributions for growing RHs. (E) Bar plot showing the average RH growth (µm) per growth period versus the average mCHERRY-RALF22$_{mature}$ microdomain spacing along the RH's longitudinal axis (µm). Data represent mean ± SEM (n=17). Asterisks indicate statistical significance (p-value; ***<0.001). (F-G) Representative mCHERRY-RALF22$_{mature}$ CW fluorescence profile for the shank (F) and tip (G) after removal of low spatial frequency components (<0.02 mCHERRY-RALF22$_{mature}$ fluorescence cycles.µm-1) by Fourier filtering. (H-I) Corresponding continuous wavelet spectra showing the cumulative prevalence of spatial frequencies for the shank (H) and tip (I) fluorescence traces. White line overlays represent the average power spectrum. (J) Average power spectra depicting the RALF22 ring spacing distributions for the tip (black) and shank (grey). Gaussian fits are depicted in green (tip) and blue (shank). (K) bar plot showing the average ring spatial frequency (mCHERRY-RALF22$_{mature}$ fluorescence cycles.µm-1) and spacing (µm) for the tip and shank. Data represent mean ± SEM (tip, n=12; shank, n=15). Asterisks represent statistical significance (p-value; **<0.01).

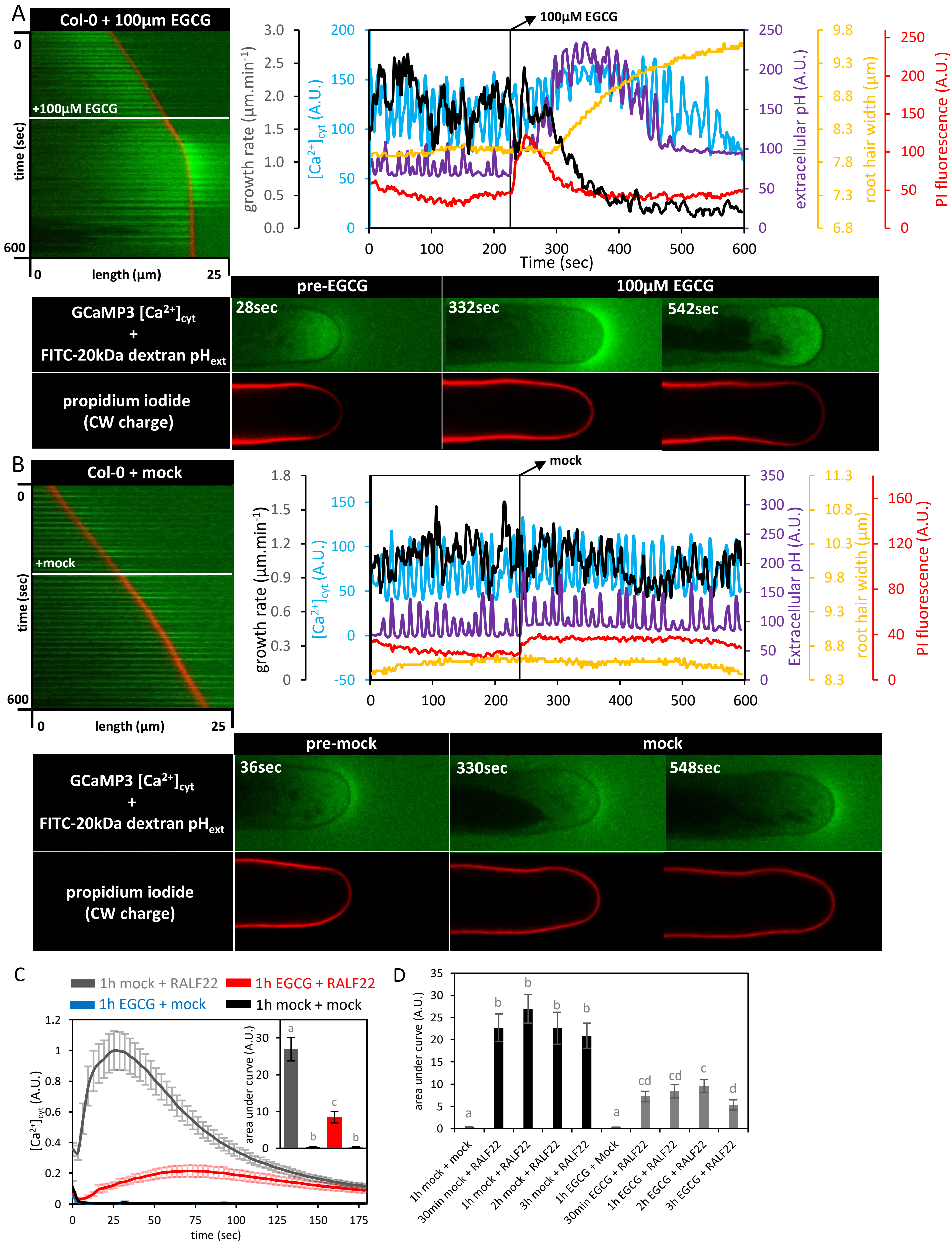


Fig. S6. pectin demethylesterification is important for root hair growth and the response to RALF22. (A-B) 10min timelapse imaging of representative Col-0 RHs being treated with 100µM EGCG (A) or growth medium (B) (cfr. movie 6). The kymographs and corresponding plots depict representative changes in the growth rate, $[Ca^{2+}]_{cyt}$ (cytosolic GCaMP3 fluorescence intensity), $pH_{ext}$ (extracellular FITC-20kDa dextran fluorescence intensity ratio), CW charge (CW PI fluorescence intensity) and tip diameter during ~4min of steady state growth and ~6min of treatment. (C-D) EGCG pre-treatment dampens the root hair-specific $[Ca^{2+}]_{cyt}$ induced by RALF22. (C) Root hair-specific RALF22- or mock-induced $[Ca^{2+}]_{cyt}$ responses of Col-0 seedlings pre-treated with EGCG (32µM) or growth medium (mock) for 1h (n=16). (D) Average $[Ca^{2+}]_{cyt}$ responses for each of the treatments, displayed as the area under the curve (A.U.). Data is shown as the mean ± SEM. Different letters represent statistical significance (α=0.05).

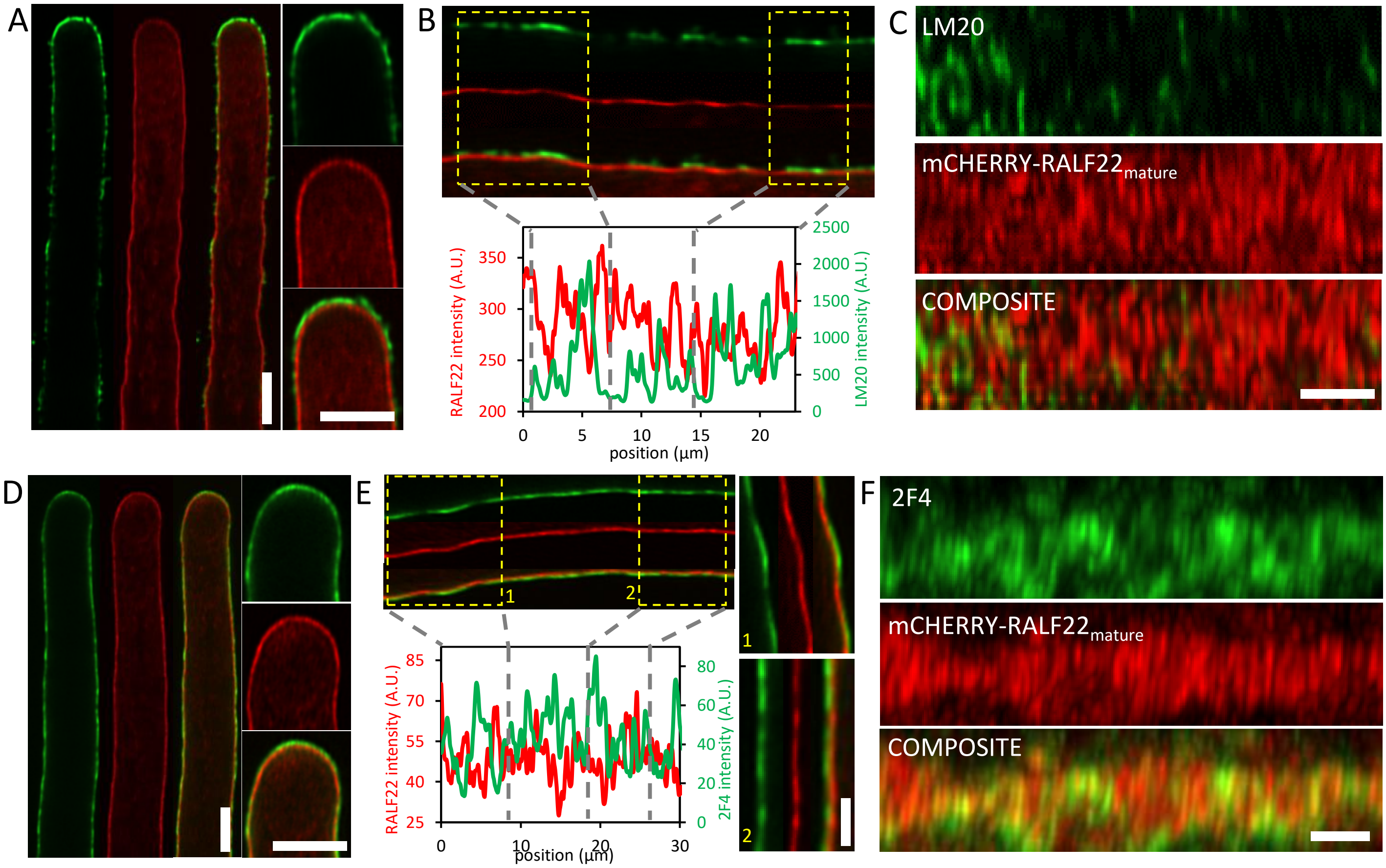


Fig. S7. localization of methylesterified and $Ca^{2+}$-crosslinked demethylesterified HG relative to mCHERRY-RALF22$_{mature}$ in the cell wall of root hairs, which were fixed during growth. (A-C) representative images showing the colocalization of LM20 (methylesterified HG) and mCHERRY-RALF22$_{mature}$ in the RH tip and shank. (A) longitudinal optical section of an LM20 (green) and mCHERRY-RALF22$_{mature}$ (red) labeled RH (scale bar=5µm) and expanding tip (scale bar=5µm). Yellow indicates colocalization. LM20 labeling is most apparent in the tip and occurs in sparse regions in the shank, where it labels an outer layer of the CW which is devoid of mCHERRY-RALF22$_{mature}$. (B) close-ups of longitudinal optical sections and corresponding fluorescence intensity traces of the labeled CW in the shank, showing LM20 labelled regions (green) that overlay intervals with overall lower mCHERRY-RALF22$_{mature}$ labeling (red). (C) lateral z-projections of the RH shank CW showing the sparsity of LM20 labelled patches relative to mCHERRY-RALF22$_{mature}$ rings (scale bar=5µm). (D-F) representative images showing the colocalization of 2F4 ($Ca^{2+}$-crosslinked demethylesterified HG) and mCHERRY-RALF22$_{mature}$ in the RH tip and shank. (D) longitudinal optical section of a 2F4 (green) and mCHERRY-RALF22$_{mature}$ (red) labeled RH (scale bar=5µm) and expanding tip (scale bar=5µm). 2F4 labels the entire RH CW in a layer which partially overlaps with mCHERRY-RALF22$_{mature}$. (E) close-ups of longitudinal optical sections and corresponding fluorescence intensity traces of the labeled CW in the shank, showing that 2F4 labelled regions (green) of lower mCHERRY-RALF22$_{mature}$ labeling (red) and vice versa (scale bar=3µm). (F) lateral z-projections of the RH shank CW showing the formation of 2F4 and mCHERRY-RALF22$_{mature}$ circumferential rings (scale bar=5µm). Regions with lower mCHERRY-RALF22$_{mature}$ abundance exhibit higher 2F4 labeling (scale bar=5µm).
